# Linking Ancient Refugia to Modern Diversity: Evidence of Multi-origin Postglacial Expansion of Sockeye Salmon on the Asian Range

**DOI:** 10.64898/2026.02.11.705351

**Authors:** Anastasia M. Khrustaleva, Ekaterina V. Ponomareva

## Abstract

Sockeye salmon (*Oncorhynchus nerka*) is a traditional object of fishery in the northern Pacific, but its island populations now face emerging threats from territories development and escalating risks of unregulated fishing. The present study aims to assess the current status of Kuril island populations associated with the biogeographical processes in the past. We analyzed diversity distribution of the D-loop sequence and three mtSNPs localized in the *Cytb* and *COI* genes in sockeye salmon populations across the North Pacific. mtDNA variants were grouped into two distinct lineages: haplogroups 10T and 13T. Their distribution suggests an Asian origin for the 10T lineage and a North American origin for the 13T haplogroup. Testing the biogeographical scenarios support recurrent postglacial expansions of North American strains into the southernmost territories of the Asian range during the last two glacial cycles. Concurrently, during the Holocene transgression, there were two centers of sockeye salmon radiation in Asia associated with the refugium in the Kamchatka River basin and a minor cryptic refugium in the Hokkaido region. We also propose an ’island bridge’ hypothesis to explain dispersal of 10T-lineage from the Kamchatka River basin into Cook Inlet and Alaska Peninsula watersheds via the Aleutian Islands.

## INTRODUCTION

Sockeye salmon is a valuable anadromous salmon species and a traditional object of fishery in the northern Pacific. Contemporary large-scale anthropogenic transformation of aquatic ecosystems, coupled with intensive commercial exploitation and global climate change, is driving substantial shifts in population structure and genetic diversity of Pacific salmon (*Oncorhynchus* spp.) across the North Pacific (Waples et al., 2008; Cline et al., 2019; Thompson et al., 2019; Koval et al., 2020; Oke et al., 2020; Crozier et al., 2021). Island ecosystems face exceptional risks under these threat conditions. Island populations typically exhibit high specialization, low genetic diversity, and small effective sizes, making them vulnerable to extinction under both short-term and chronic environmental stressors. Their extreme sensitivity to external perturbations requires targeted conservation strategies, including flexible fishery management, as most sockeye salmon populations in the Russian Far East islands remain under exploitation. Research aimed at maintenance of commercial fish biodiversity and the sustainable stock management is the main focus of applied ichthyological science. At the same time, island populations may become an essential model for understanding microevolutionary dynamics and reconstructing the evolutionary history of the species on the Asian Pacific coast.

Currently, sockeye salmon populations of the Kuril and Commander Islands inhabit sparsely populated regions that have experienced minimal anthropogenic impact. However, intensive economic development of these territories, infrastructure expansion in the river basins, unregulated fishery or inflexible exploitation management could cause significant damage to these stocks. The South Kuril Islands populations, located at the southern range margin of Asian sockeye salmon, are particularly vulnerable to global warming. Consequently, assessing their current conservation status represents an important research interest within this study.

The distribution and population structure of the species across its range was evidently linked to climate oscillations during the Middle and Upper Pleistocene (Shafer et al., 2010). According to existing hypotheses (Varnavskaya et al., 1994; Wood et al., 1994; Bickham et al., 1995; Beacham et al., 2006a), contemporary sockeye salmon populations in the rivers and lakes of the USA and Canada originated from two major glacial refugia: a northern refugium − Beringia (encompassing the exposed continental shelves of the northern seas, Chukotka, the Anadyr River basin, and Alaska up to the Mackenzie River in Canada) and a southern refugium − Cascadia (situated south of the Cordilleran Ice Sheet, within the Columbia River basin). Additionally, some authors suggest the presence of small mountain refugia for the sockeye salmon on the Queen Charlotte Islands and Vancouver Island, which became exposed during the glaciation due to lowered sea levels (Wood et al., 1994). More recent studies have shown that there can be some refugia complexity (refugia within refugia) and there are strong evidences for cryptic refugia in some areas on the East Pacific coast (Shafer et al., 2010). For instance a distinct group of kokanee from the upper Columbia River was identified using WGS data that might be a glacial relict isolated during the Last Glacial Maximum (or LGM, ∼26.5–19 thousand years ago (ka)) in large periglacial lakes that connected the Columbia and Fraser Rivers (Christensen et al., 2020).

In contrast, the phylogeography of Asian sockeye salmon remains poorly understood, with most studies restricted to specific regions of the species distribution (Bickham et al., 1995; Churikov et al., 2001; Brykov et al., 2005; Kondzela & Gharrett, 2007; Bachevskaya et al., 2015; Khrustaleva, 2016; Khrustaleva et al., 2020)The hypothesis has been repeatedly originated that most populations of Asian sockeye salmon are relatively young, and their age does not exceed 10-12 thousand years, since its modern range largely coincides with the supposed area of the last Wisconsin (Wurm, according to the interregional correlation scheme) glaciation (∼75–11 ka, deglaciation ∼16.9−12.68 ka) in the North Pacific (Chereshnev, 1998; Brykov et al., 2005). Nevertheless, geological and genetic evidences have led to an assumption of the existence of a large glacial refugium during LGM in the Central Kamchatka Depression (Varnavskaya et al., 1994; Brykov et al., 2005; Khrustaleva et al., 2020). There are also speculations about the existence of a cryptic refugium located south of the ice-covered Kamchatka and northern Kuril Islands, in the area of Primorye, Sakhalin and Hokkaido Islands (Khrustaleva et al., 2020). The present study aims to evaluate these two hypotheses using a phylogeographic approach based on diversity distribution of two mtDNA markers (complete D-loop sequence and three mtSNPs) across the Asian range. Moreover in order to develop conservation strategies for the unique genofond of island sockeye salmon populations, we will attempt to characterize the current genetic diversity of island forms, in particular the populations of sockeye salmon of the South Kuril Islands, and consider possible scenarios for the paleocolonization of the southernmost territories of the Asian range. Data on mtSNP variants for the entire range of sockeye salmon will allow us to draw conclusions about the origin of the population structure of the species and possible expansion patterns of sockeye salmon after the LGM.

## METHODS

### Sample collection, DNA extraction and sequencing

A total of 280 sockeye salmon specimens from 28 samples collected in 2003-2015 in watersheds of the Chukotka Peninsula (Chukotka), the Kamchatka Peninsula (Kamchatka), the mainland coast of the Sea of Okhotsk, the Kuril and Commander Islands were analyzed (Fig. 1a, Table 1). Pectoral fin fragments fixed in 96% ethanol were used for genetic analysis. Total genomic DNA extraction from the fin tissue was conducted by DNeasy Blood & Tissue Kit (QIAGEN) according to the manufacturer’s protocol. HN20 and Tpro2 primers (Brunner et al., 2001) were used for complete mtDNA control region (D-loop) amplification (for 189 specimens). For 91 fish, a 702 bp fragment (336–1037 bp in the mitochondrial genome) from the D-loop sequence adjacent to tRNA1 was amplified using self-designed primers (primer sequences: F: TGTAGTAAGAACCGACCAACGAT and R: ACTTTCTAGGGTCCGTCTTAACAGC). PCR was conducted in total of 20 μL volume containing 2 μL of 10Х AmpliEN buffer, Mg2+ 2.0 mM, 0.125 mM dNTP’s, 1U BioHYTaq DNA polymerase (DIALAT, Russia), 5 pmol of each primer, 1.5 μL template DNA. The amplification was performed in the following mode: I – 95°С 5 min., then 35 cycles II – 94°С 10 sec., 20 sec. Тa, 56°С, 72°С 1 min., III – final elongation 72°С – 10 min. Horizontal electrophoresis in 1.5% agarose gel was carried out for PCR product control. The remaining amplicon was purified by precipitation following the protocol (CCU“Genome,” 2011). Forward and reverse Sanger sequencing of the 702 bp D-loop fragment were performed on an automatic sequencer (ABI PRISM 310 Genetic Analyzer, Applied Biosystems, USA) using the ABI PRISM BigDye Terminator version 3.1 reagent kit and the same self-designed primers. Full D-loop sequencing was conducted in the CCU “Genome”. Overall 108 new D-loop sequences of Asian sockeye salmon were generated under this study. In addition, the data on 172 entire D-loop sequences were used (Khrustaleva et al., 2020).

**Fig. 1.**
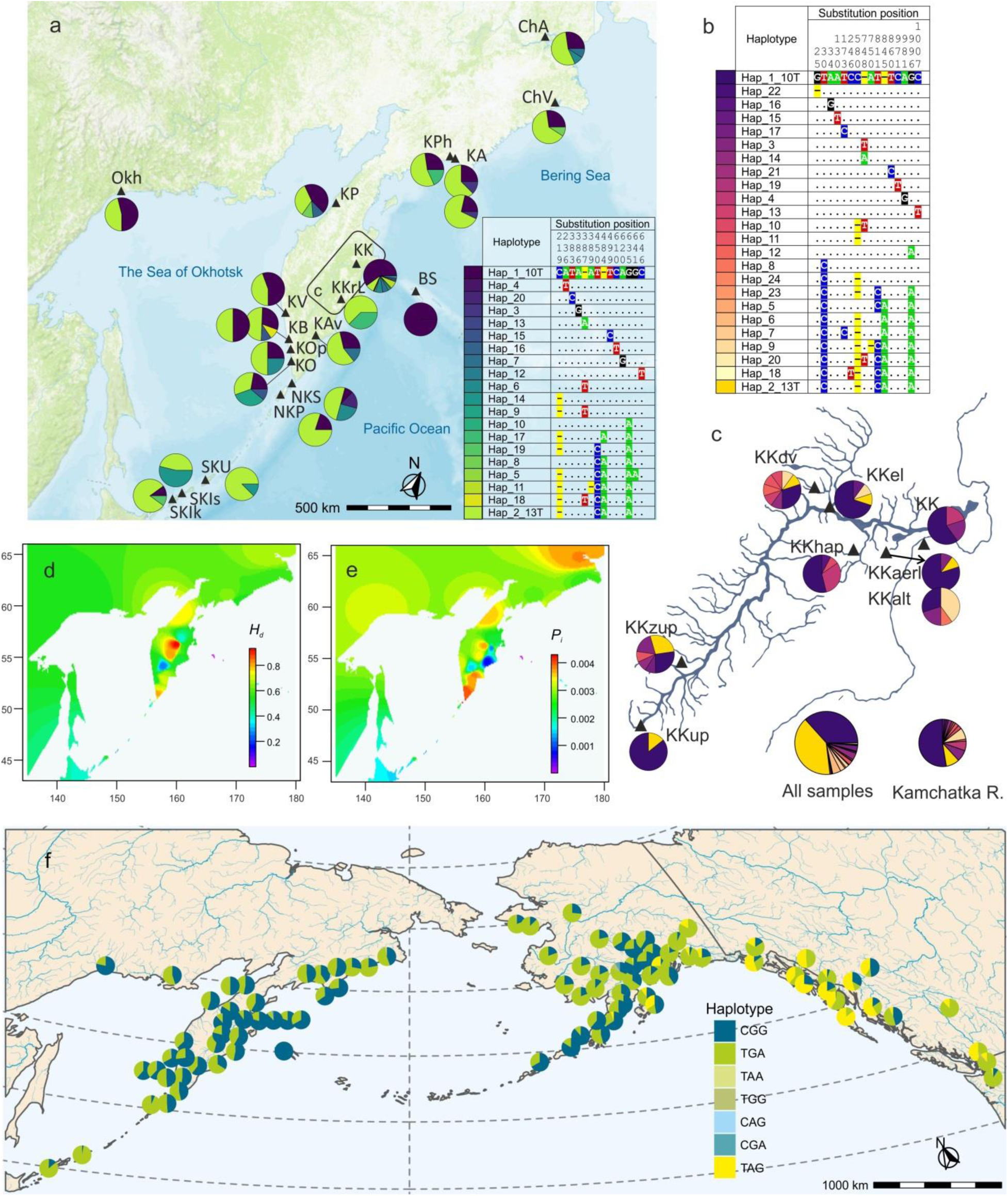
Map of sampling locations on the Asian Pacific coast and the short−length set haplotypes frequencies in the samples **(a)**, a key to the charts with the D−loop region variable cites is given in the table. **(b, c)** The same for the long−length set haplotype frequencies in the Kamchatka River drainage. Spatial distribution of genetic diversity estimates *H_e_* **(d)** and *P_i_* **(e)** across the Asian part of the sockeye salmon range. **(f)** Geographical distribution of mtSNP haplotypes across the entire range of sockeye salmon

**Table. 1.**
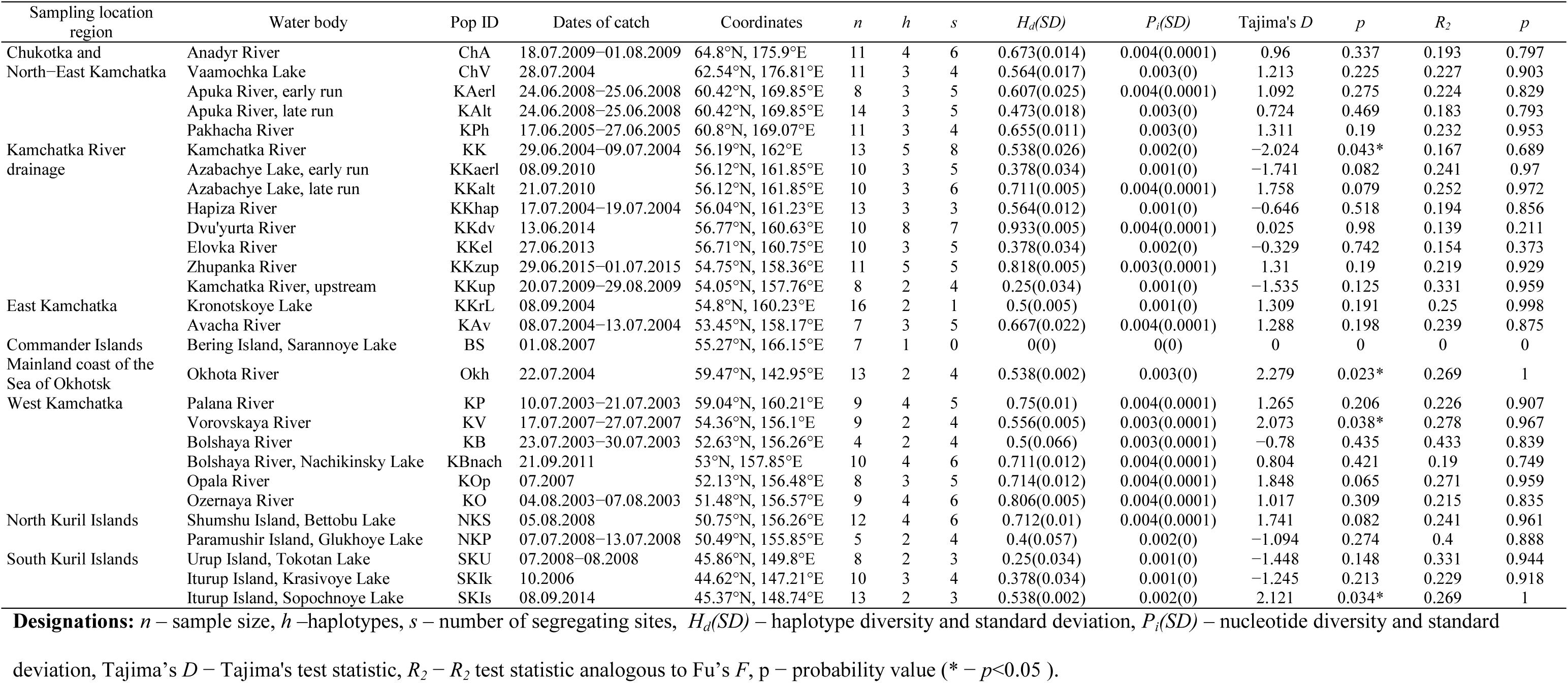
Samples characteristics, regions and locations, population IDs, genetic diversity indices and results of mutational−drift equilibrium tests for sockeye salmon based on the mtDNA D−loop sequences.

### Primary processing and multiple alignments of sequences

Primary processing of nucleotide sequences was conducted in Geneious 6.0.5 (Biomatters Ltd.). Final multiple alignment was performed Jalview 2.11.4.1 (Waterhouse et al., 2009). The two sets of data were obtained: the first set of the aligned D-loop fragments of 702 bp length, which were trimmed up to 674 bp according to the length of the shortest sequence in the array, included 280 sequences, whereas the second one, included 189 sequences of the complete mtDNA control region, 1035 bp in length. Since most phylogenetic tools ignore indels, resulting in significant loss of information, all indels were artificially replaced by transversions in the final alignments. FaBox (Villesen, 2007) was used for converting the allinments into haplotypes sets and input files preparation.

### Genetic diversity

The set of short D-loop sequences (674 bp long) was used to calculate the diversity statistics such as number of haplotypes (*h*) and segregating sites (*s*), haplotype diversity (*H_d_*), nucleotide diversity (*P_i_*) in the pegas R package (Paradis & Barrett, 2010). To visualize the spatial pattern of the sequences diversity across landscapes, interpolation surfaces of nucleotide diversity (*P_i_*) and haplotype diversity (*H_d_*) estimates were generated with R function GDivPAL from the SPADS toolbox (Dellicour & Mardulyn, 2014). Calculation and interpolation of nucleotide diversity (*P_i_*) were performed using Sliding Window Algorithm with SPADS R functions slidingWindowPi. The resulting rasters were visualized with the maps R library (Becker et al., 2022). To detect deviation from mutational-drift equilibrium in the populations, the Tajima’s *D* and *R_2_*, analogous to Fu’s *F*, but more appropriate for small sample sizes (Ramos-Onsins & Rozas, 2002), neutrality tests were calculated in pegas. Significantly negative values for both statistics are generally taken to indicate a sudden population growth (expansion) in the past and/or positive selection, while significantly positive estimates indicate a recent population decline (bottleneck), population fragmentation, and/or balancing selection. Because of the functional neutrality of mtDNA, statistically significant tests are usually considered in the context of demographic events. Non-metric multidimensional scaling (MDS) of *F_ST_* values between samples calculated by Tajima and Nei’s method in Arlequin v. 3.5. (Excoffier & Lischer, 2010) was performed to visualize genetic similarity.

### Phylogeographic, phylogenetic and historical demographic analyses

MST (minimum spanning tree) genealogies for the both sets of mtDNA haplotypes (short and long) were constructed using the pegas package. Phylogenetic relationships between haplotypes were inferred using maximum likelihood (ML) trees reconstruction in IQ-TREE 2.0.3 (Nguyen et al., 2015) with an Unequal transition/transversion rates and unequal base frequencies model (Hasegawa et al., 1985) using empirical base frequencies and allowing for a proportion of invariable sites (HKY+F+I) as a best-fit model by all criterions (AIC, AICc, BIC) calculated in ModelFinder modul implemented in IQ-TREE. As an out-group *Oncorhynchus keta* sequence (LC588924.1) was used. Node support values were estimated from 1000 bootstrap replicates.

Haplotypes divergence time was estimated by constructing a time-calibrated phylogeny as implemented in BEAST 2.6.2 (Bouckaert et al., 2019). The only complete mtDNA control region haplotypes dataset was used to reconstruct a phylogenetic tree using two calibration points: the first one was the fossil † *Oncorhynchus keta* (minimum age − 4.8 Ma, maximum 95% CI age − 40.55 Ma) (Smith, 1992) (LC588924.1 and NC_010959.1 for *Oncorhynchus gorbuscha* were used as an out-group), while the second was a geological event, a volcanic eruption that led to the isolation of the population of Kronotsky Lake, at the beginning of the Holocene (about 10 thousand years ago) (Zhivotovsky et al., 2019). The analyses were run using a strict clock algorithm, the site model was chosen as HKY+F+I, and the mutation rate was set as 0.01 substitutions site-1, myr-1(Bermingham et al., 1997; Stepien et al., 1997; Grant, 2015). The calibrated Yule model was used as a tree prior. A lognormal distribution was assumed for the fossil age (with the following parameters: *µ* = 1.57 and *σ* = 0.7 resulting in median of 4.8 Ma, and 97.5%−quantile of 19.0 Ma) and a normal distribution for the geological event (*m* = 0.01, *σ* = 0.001 resulting in median of 10,000 years, and 97.5%−quantile of 8,000-12,000 years). All other model parameters used default priors. MCMC was run for 50 million generations, with samping every 1000th generation and removing the initial 10% of samples as burn-in. The convergence of the chain to the stationary distribution was confirmed by inspection of MCMC samples using Tracer v1.7.1 (Rambaut et al., 2018). Results from the independent runs were compiled with 10% burn-in using LogCombiner v2.6.2 (Bouckaert et al., 2019) and then summarized for the maximum clade credibility (MCC) tree in TreeAnnotator v2.6.2 (Bouckaert et al., 2019). Tree visualization was performed in FigTree 1.4.4 (Rambaut, 2018), as well as ggtree (Xu et al., 2022) and deeptime (Gearty, 2024) R packages.

To examine the possible occurrence of a historical demographic expansion of each phylogenetic lineage (haplogroup), we performed mismatch distribution analysis (Rogers & Harpending, 1992), which was implemented in Arlequin 3.5 (Excoffier & Lischer, 2010). The fitness of the observed population data to the models of sudden population expansion and spatial expansion were tested by calculating the sum of the squared deviation (SSD), the models were accepted if *p* > 0.05. The demographic history of the sockeye salmon in Asia was also examined by Bayesian skyline plot (BSP) analysis (Drummond et al., 2005), implemented in BEAST 2.6.2. We performed runs with a MCMC chain length of 10 million generations, sampling every 1000th generation and removing the initial 10% of samples as burn-in. The substitution model used was the HKY+F+I, and the time to expansion was estimated using the mutation rate 0.01 substitutions site-1, myr-1. Data visualization was performed in R using ggplot2 library.

### Biogeographic scenario testing

To test competing hypotheses about the biogeographic pathways leading to the formation of the Kuril Islands populations from Kamchatka and North American stocks, an approximate Bayesian computation (ABC) approach within a coalescent framework implemented in DIYABC 2.1.0 (Collin et al., 2021) was used. Four biogeographic scenarios represented by their historical models (with a tree-like graphic representation) were tested. Prior settings of the ABC analyses are presented in Supplementary Table S1. The average generation time for sockeye salmon was taken as 4 years. The prior values were used with a uniform distribution for all parameters. A total of 4,000,000 simulated datasets were calculated. Pre-evaluations of model prior combinations in the ABC inference were used to assess whether the prior settings were assigned correctly. Posterior probabilities of the four biogeographic scenarios were calculated using direct and logistic approaches. The goodness-of-fit of each scenario was assessed via a principal component analysis (PCA) of the prior and posterior parameters, performed using the “model check” option in DIYABC.

### mtSNP data

To trace the distribution of mtDNA variants across the entire range of sockeye salmon, we used both the own data on the frequencies of three mtSNPs genotyped by TaqMan-PCR (Seeb et al., 2009) and the open data on the same set of loci in a wide range of Asian and American populations (Habicht et al., 2010),(Habicht et al., 2010) as well as the data on mass mtDNA haplotypes ratio for populations from the Tauysky Bay (Ola River, *n* = 48) and Karaginsky Gulf (Khaylulya River, *n* = 48) (Bachevskaya et al., 2015)(Bachevskaya et al., 2015) (Supplementary Table S2, S3). The *One_Cytb_17* (A/G transition) and *One_Cytb_26* (C/T transition) loci are located in the *Cytb* gene, at positions 16162 and 16168 of sockeye salmon mtDNA (GenBank EF055889.1), and the *One_CO1* (C/T transition) is located in the *COI* gene encoding cytochrome oxidase subunit I, at position 7061. All three SNPs are represented by synonymous substitutions in the third position of the codon and were concatenated into a combined locus *CO_Cytb* (*One_CO1*−*One_Cytb_17*−*One_Cytb_26*) for subsequent analysis. In total, the mtSNP database included 40 Asian and 55 North American sockeye salmon populations from the largest breeding watersheds on both coasts of the North Pacific Ocean. Visualization of the geographic distribution of sockeye salmon *CO_Cytb* variants across the range was performed using the sf (Pebesma & Bivand, 2023) and ggplot2 (Wickham, 2009) R libraries.

## RESULTS

### D-loop sequence diversity and haplotypes distribution

After multiple alignment of the 280 D-loop sequences the two datasets were generated: the "short-length" set of the 674 bp D-loop fragments included all 280 sequences, and the "long-length" set of the complete mtDNA control region sequences, 1035 bp in length, contained 189 sequences (Khrustaleva et al., 2020). Overall 20 "short-length" haplotypes were discovered for the first time and 24 "long-length" haplotypes (including a new haplotype (Hap_20) found in Nachikinsky Lake sample(Khrustaleva et al., 2020)) were detected in sockeye salmon populations on the West Pacific Coast (Fig. 1a, b, c). Two mass haplotypes (Hap_1_10T and Hap_2_13T) were identified in the entire dataset; they were observed in most of the samples and differed by two/tree ("short-length"/"long-length") substitutions and two indels (Fig. 1a, b). According to genealogy of Asian sockeye salmon D-loop haplotypes all sequence variants were distributed among two haplogroups/phylogroups/phylogenetic lineages 10T and 13T (Fig. 2) with the central haplotypes Hap_1_10T and Hap_2_13T. Both lineages formed star-shaped topology; minor/rare haplotypes differed from central ones in one or two substitutions. The haplogroup 10T was slightly more numerous, mainly due to the greater haplotypes diversity.

**Fig. 2.**
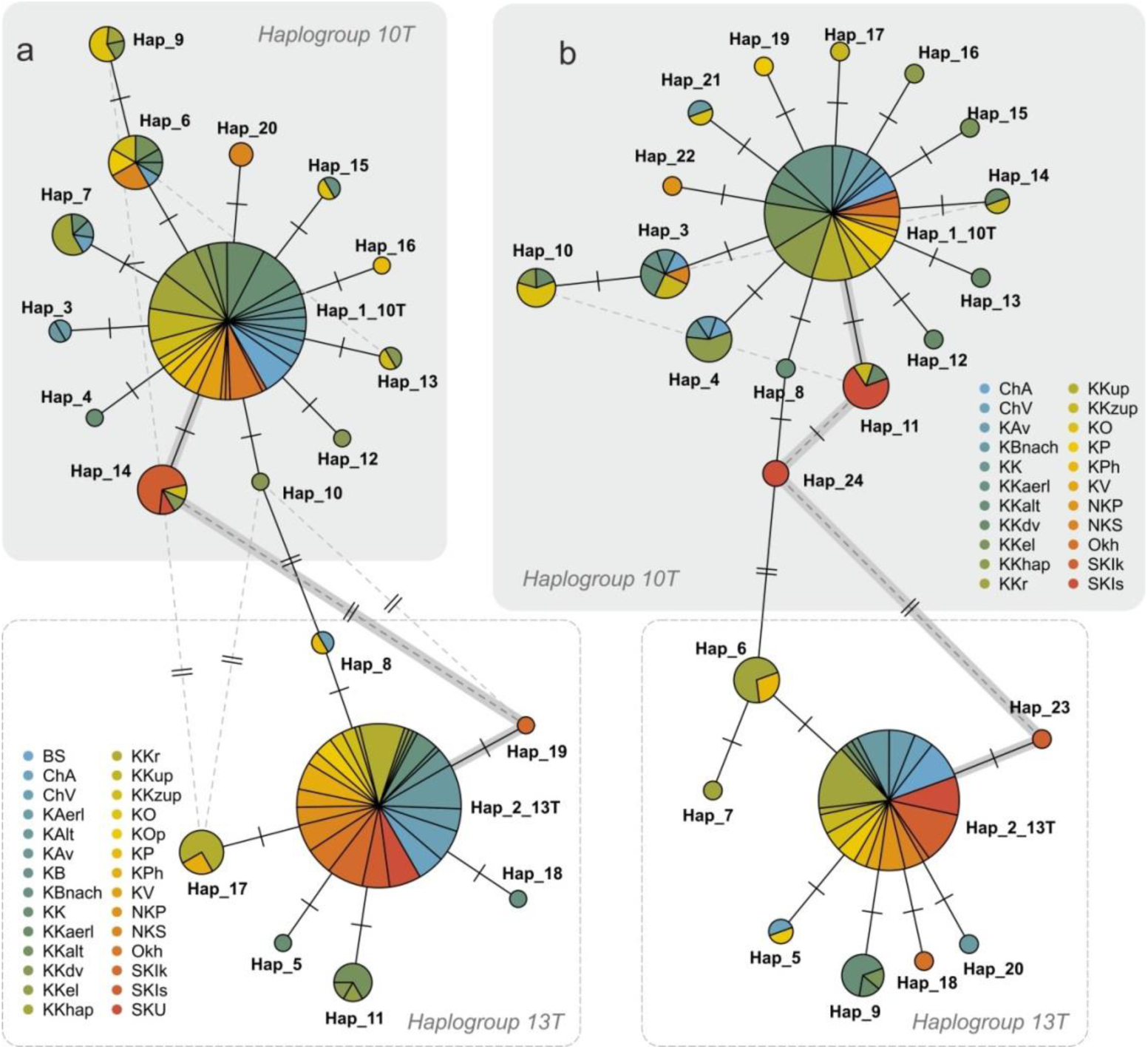
Genealogical networks of sockeye salmon D−loop haplotypes, built on the base of the minimum number of nucleotide substitutions (MST−tree): **(a)** – for short−length set, **(b)** – for long−length set. The size of the circles and the area of the sectors are proportional to the haplotypes frequency in the corresponding samples. The solid grey lines are the most probable paths of haplotype transformation

Further results will be presented for the "short-length" sequences set, as more representative data are available for these sequences, whereas the "long-length" set has been partially characterized in (Khrustaleva et al., 2020). The spatial distribution of haplotypes along the coast of the Sea of Okhotsk and the Pacific coast was somewhat different: if along the Sea of Okhotsk coast both haplotypes were found in approximately equal proportions, then along the Pacific coast Hap_2_13T predominated, with the exception of the populations of the Kamchatka River basin and Lake Sarannoye (Bering Island) (Fig. 1а, c).

Data on sockeye salmon D-loop sequences diversity on the Asian range are summarized in Table 1 and Figures 1d, 1e. The greatest haplotype diversity is observed in the Kamchatka River basin (Fig. 1c), in addition, high estimates of haplotype and nucleotide diversity were characteristic of populations from the largest sockeye salmon reproduction watersheds on the western coast of the Pacific Ocean.

### mtSNP distribution across the entire sockeye salmon range

Six haplotype variants of the *CO_Cytb* locus (mtSNP haplotypes) were detected in the Asian samples, two of them (CGG and TGA) were mass, while the others were singly present. The correlation of the two main mtSNP haplotypes and the variants of the nucleotide sequences of the mtDNA control region in all cases where the same samples were genotyped by both markers allowed us to establish the congruence of the common haplotypes of Asian sockeye salmon in the overwhelming majority of comparisons: in more than 90% of cases, the CGG SNP haplotype corresponded to the Hap_1_10T haplotype, and TGA to the Hap_2_13T. The high degree of similarity in the variability of different mtDNA regions in sockeye salmon is evidenced by previously obtained experimental data (Churikov et al., 2001). The revealed discrepancies can be explained by both methodological (genotyping errors, sample identification errors) and objective reasons related to haplotype diversity within haplogroups. Thus, as a first approximation, we can consider the mass SNP haplotypes (CGG & TGA) identical to two phylogenetic lines of sockeye salmon from the northern part of the Pacific Ocean (respectively, Haplogroup 10T & Haplogroup 13T).

The distribution of the frequencies of the main SNP haplotypes across the sockeye salmon range shows that both haplotype variants are found in most populations (Fig. 1f); only the sample from Sarannoye Lake (Bering Island) was represented by a single CGG haplotype, probably due to the founder effect during the colonization of the island’s watersheds by migrants from the Kamchatka River basin. Haplogroup 13T was predominant in the Asian sockeye salmon populations of northeastern Kamchatka and Chukotka and the northernmost populations of the American coast (Norton Sound, Yukon and Kuskokwim rivers, as well as in the watersheds of the North-East coast of Bristol Bay). However, in the Kamchatka River basin, the watersheds of the Commander and Aleutian Islands, and the lake-river systems of the Alaska Peninsula and Cook Inlet, haplogroup 10T was more numerous. In the Kodiak region and the northern and eastern coasts of Alaska Bay (except for the Nass and Skeena River basins), haplogroup 13T was again numerically predominant, while in samples from this region, the SNP haplotype TAG, endemic for this region, was encountered with a fairly high frequency. The origin of TAG haplotype is most likely associated with the South American Cascadia refugium. On the Asian coast in the southernmost part of the range (the Kuril Islands except for Shumshu Island), the 13T line was again widespread.

### Population structure and historical demography

According to the multidimensional scaling of F_ST_ values calculated for the "short-length" dataset, samples from distinct regions formed more or less dense clusters with the exception of the South Kuril Islands populations and the sample of kokanee from the Kronotsky Lake (Fig. 3a).

**Fig. 3.**
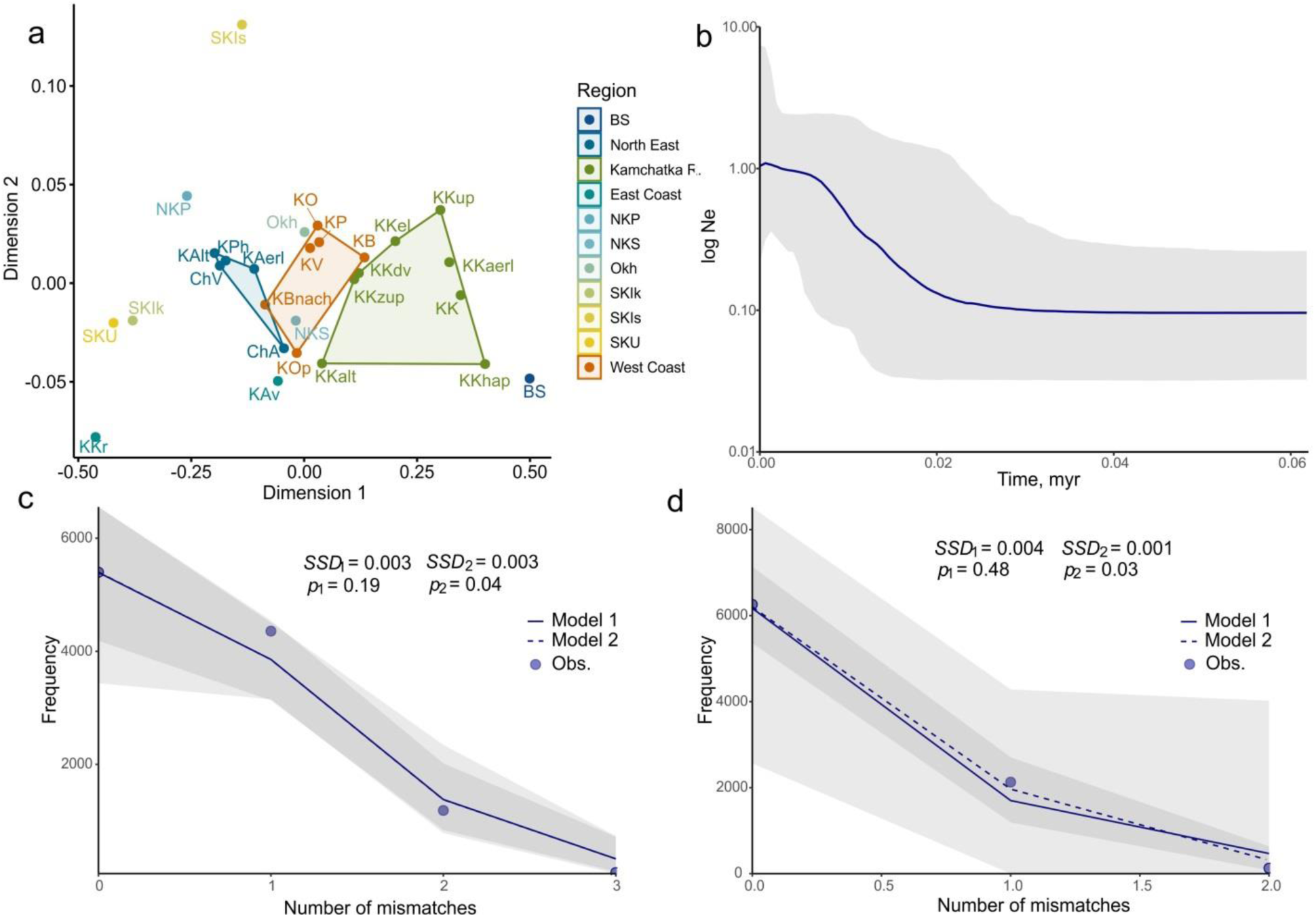
Multidimensional scaling plot based on the pairwise *F_ST_*’s estimates for the short−length set of the D−loop sequences **(a)**, regions are marked by different colors, some populations are combined into polygons according to region, some are outliers. Bayesian skyline plot for the Asian sockeye salmon **(b)**, the central bold line represents the median value for the relative effective population size, and the solid area denotes the 95% upper and lower credible limits. Mismatch distribution for the two phylogenetic lineages of sockeye salmon: **(c)** – haplogroup 10T, **(d)** – haplogroup 13T. The solid lines represent the expected distributions under the sudden expansion model (Model 1), the dashed line – the expected distributions under the spatial expansion model (Model 2), dots – frequencies of the observed pairwise differences

*R_2_* tests did not reveal significant deviations from the mutation-drift equilibrium in any of the samples (Table 1). At the same time, the *D*-test results revealed traces of a recent decline in population size (bottleneck) in three samples, and for the Okhota River population there is documented evidence of a significant decrease in *Ne* in the middle of the last century (Nikulin, 1975; Marchenko, 2022). In the sample from the mouth of the Kamchatka River, an excess of new mutations (significantly negative *D*) was revealed, most likely due to a sharp increase in population size (expansion) in the relatively recent past.

Skewed unimodal mismatch distributions were obtained for two phylogenetic lineages of the Asian sockeye salmon (Fig. 3c, d). Such distributions are generally associated with a recent sudden expansion or bottleneck (Jenkins et al., 2018). The observed data fitted well the expected distribution under the model of spatial expansion (accepts as the null hypothesis a one-time expansion of the range from one deme/population, followed by an increase in the number and the formation of new demes/populations). As can be seen from the plot (Fig. 3c, d) the sudden expansion model seems to be less appropriate for both lineages (none of the sums of squared deviations (SSD) of the mismatch distribution was significant). In order to assess the time of the range expansion in Asia a Bayesian skyline plot (BSP) was reconstructed (Fig. 3b). The BSP shows that the sockeye salmon Ne rapidly increased significantly after the last Wisconsin glaciation maximum (∼26.5−19 ka) in the North Pacific (Braitseva & Evteeva, 1968; Velichko & Faustova, 1989).

### Phylogenetic reconstructions

The structure of the ML trees constructed for the two sets of sequences agreed well with the genealogy of haplotypes: there were distinguished two clades of haplotypes corresponding to two haplogroups, within the clades a polytomy of sequences differing by only one substitution was observed (Fig. 4a, b). Based on the obtained trees topology, it can be concluded that the most probable path of divergence of the two phylogenetic lineages of sockeye salmon is the variant where the intermediate haplotypes was Hap_19 and Hap_14 for the "short-length" set and Hap_23, Hap_24 and Hap_11 for the "long-length" set. Nevertheless, the bootstrap support for the obtained topologies was relatively low.

**Fig. 4.**
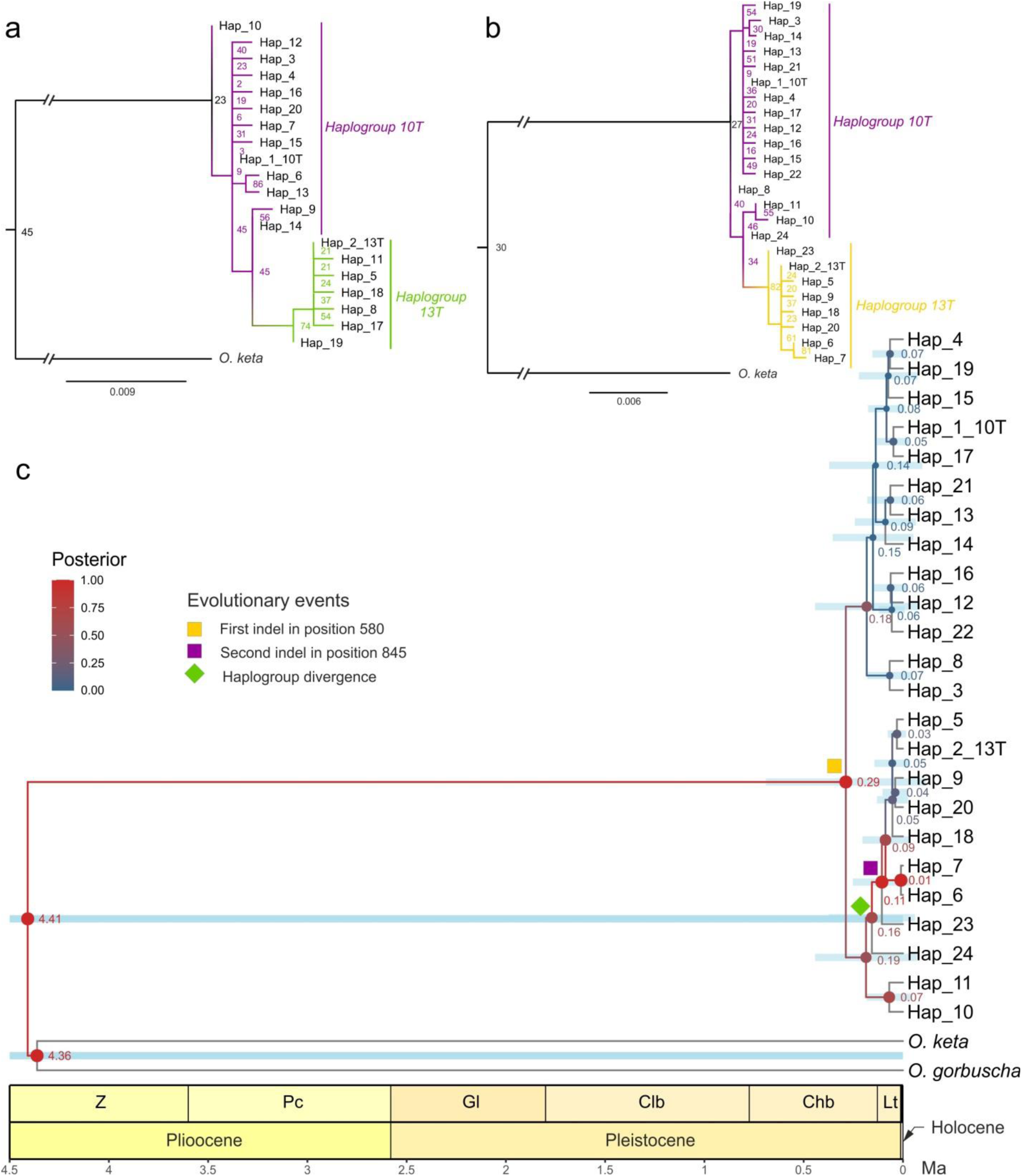
Maximum likelihood trees for short−length set **(a)** and for long−length set **(b)** of the D−loop sequences, numbers in the nodes are bootstrap indices. Calibrated Bayesian phylogenetic tree **(c)** for the entire D−loop haplotypes of Asian sockeye salmon. Values on nodes indicate node ages (Ma), colored and different−sized dots represent posterior probabilities, and light blue bars are 95% CI for the ages

The Bayesian tree for the full D-loop sequence of sockeye salmon, dated using paleontological data and geological event (volcanic eruption and damming of the Paleo-Kronotsky river by the lava dam approximately 10 ka), was congruent with our ML trees and supported our conclusions about the most probable pattern of haplotype genealogy (Fig. 4c). Posterior probabilities were high only for basal nodes (0.65 and higher), within the haplogroups the branchings had no statistical support (probability below 0.2).

### Biogeographic scenario testing

Biogeographic scenarios of the Kuril Islands colonization were reconstructed on the basis of alternative hypotheses about the sockeye salmon populations origin in these territories (Fig. 5a). The following probable pathways of sockeye salmon invasion into the watersheds of the Kuril Ridge in the Pleistocene-Holocene were considered:

**Fig. 5.**
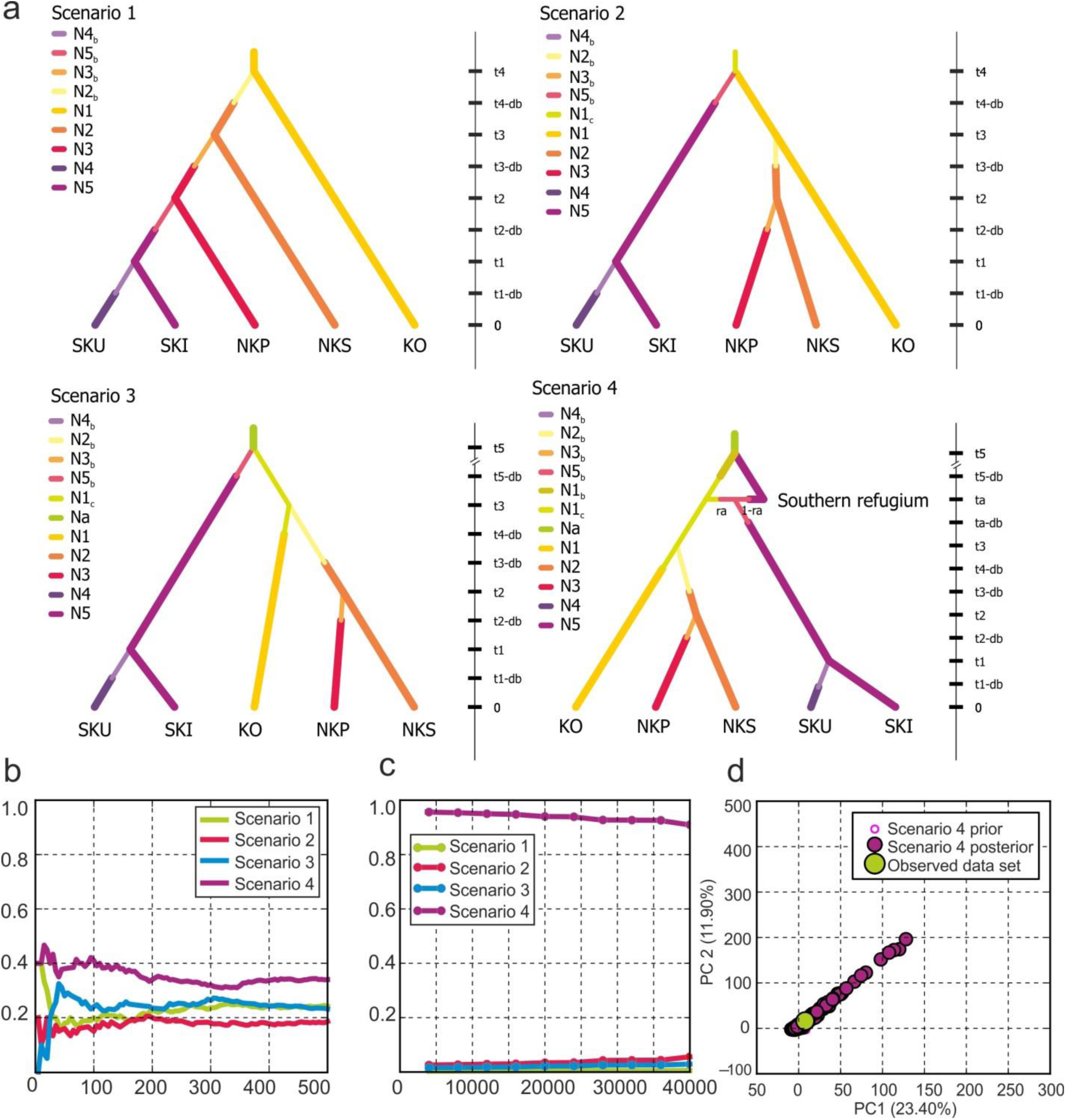
The results of biogeographic scenario testing to identify the pathways leading to the formation of the Kuril Islands populations. **(a)** Fore biogeographic scenarios designed and tested in ABC analysis. The most likely scenario with the highest posterior probability identified by using a direct approach **(b)** and weighted logistic regression **(c)**. Scenario 4 was the most probable, while possibilities for other scenarios were negligible. **(d)** Model checking results showing the agreement between simulated and observed datasets by applying a PCA to the best−supported scenario 4 based on 10,000 simulations

Scenario 1 − a classic invasion history from an ancestral population (the nearest large population of the Kuril Lake, Ozernaya River basin) in Holocene;

Scenario 2 − parallel colonization by two strains in Holocene: the Northern Kuril Islands were populated sequentially by migrants from the Kuril Lake or simultaneously with them, while the colonization of the southern islands occurred by a parallel migration stream from the Beringia and Kamchatka refugia in the late Pleistocene − early Holocene;

Scenario 3 – the same as Scenario 2, but it is assumed that the invasion in the southern islands took place much earlier − in the Middle Pleistocene during the Sangamonian interglacial (Eemian; ∼130–80 ka);

Scenario 4 – the same as Scenario 3, but slightly more complex, since it implies recurrent unidirectional introgression of genes from northern populations into the ancestral population of the southern territories (Hokkaido Island) in the Late Pleistocene during the repeated expansion of the species in the Asian part of the range, where a fraction of genes (ra) from the northern population with an effective population size of N1_b_ is introduced into the southern population (Hokkaido/Southern refugium) at time t_5_.

According to the results of the ABC analysis, the most probable scenario was the most complex scenario 4, with the highest posterior probability estimates *p*(95% *CI*) = 0.956(0.916–1)/0.41(0−0.84) (for logistic/direct approach), while the other scenarios were poorly supported (scenario 1: *p*(95% *CI*) = 0.005(0−0.6449)/0.24(0−0.62); scenario 2: 0.056(0−1)/0.2(0−0.55); scenario 3: 0.029(0−1)/0.28(0−0.67)) (Fig. 5b, c). The type I and mean type II error values for the scenario 4 were 0.195 and 0.198(0.146–0.29) for logistic regression respectively, and 0.187 and 0.12(0.08–0.18) for direct approach. The scenario and the parameters priors matched the data summarized by the summary statistics so it can be concluded that the chosen model correctly explained the observed dataset (Fig. 5d).

## DISCUSSION

### Patterns of Haplotypes and Haplogroups Distribution across the North Pacific

The geographic distribution of mDNA strains along the northern Pacific coast and the presence of both haplogroups in most Asian populations and three mtDNA lineages in North American populations confirm our earlier hypothesis that the entire range of sockeye salmon is a complex mosaic zone of secondary contacts of three large ancient allopatric gene pools isolated during the late Pleistocene glaciations in refugia on the territory of the contemporary sockeye salmon range. The star-shaped topology of the D-loop haplotype network indicates a significant increase in the diversity of Asian phylogroups due to the rapid explosive expansion of the modern range of the species after its decline during Pleistocene climate oscillations. The hypothesis of the spatial expansion of sockeye salmon on the Asian coast during the Holocene transgression is supported by the results of the mismatch distribution and BSP analysis.

Spatial distribution of mtDNA haplotype frequencies indicates a widespread distribution of 13T phylogroup throughout the entire range of sockeye salmon: on both continents 13T haplogroup was predominant in the northernmost and southernmost populations. This fact and the limited haplotype diversity of 13T phylogenetic lineage in the Asian part of the range may be an argument in favor of the hypothesis of its adventive American origin and recurrent large-scale expansions of American sockeye salmon populations on the Asian coast during the Pleistocene-Holocene interstadials. On the contrary, the high haplotype diversity identified in the 10T lineage on the Asian coast indicates its connection with the Asian coast of the Pacific Ocean: either aboriginal origin or its long-term existence in this territory.

### Spatial Distribution of Genetic Diversity in the Asian Range and Putative Refugia

It has been repeatedly suggested by different authors (Beacham et al., 2006b; Varnavskaya, 2006; Bugaev & Kirichenko, 2008; Khrustaleva et al., 2020) that during the Upper Pleistocene climate fluctuations, one of the large refugia in the Asian part of the sockeye salmon range was located in the Kamchatka River paleobasin. According to our data, extremely high estimates of haplotype diversity were found in samples from the Kamchatka River basin, where in addition to the two main haplotypes, 10 more derivative haplotypes were found, while in other watersheds only 1−3, maximum fore haplotypes were present in total. Obviously, in areas not subject to glaciation, genetic diversity should be significantly higher than the background (Hewitt, 1996). The existence of a refugium in the Kamchatka River basin is also confirmed by the presence here of a relatively rich freshwater malacofauna and the habitat of a typical representative of the freshwater Far Eastern ichthyofauna - the Kamchatka grayling (*Thymallus arcticus mertensi*), which penetrated the Kamchatka River along the connected watersheds of the Central Kamchatka Depression and survived here during the GLM (Kurenkov, 1965; Chereshnev, 1998; Prozorova & Shedko, 2003). However, there is no evidence that populations of other Pacific salmon species persisted in the Kamchatka refugium (McCusker et al., 2000; Shpigalskaya et al., 2008, 2009; Savin et al., 2009; Bachevskaya et al., 2011, 2017, 2021; Zelenina et al., 2020). It is likely that sockeye salmon, the only species capable of transition to a long-term freshwater lifestyle as a resident form, kokanee, was able to survive long-term isolation in the lake, while the Paleo-Kamchatka River discharge and oceanic outlet for anadromous fishes during Pleistocene maxima were most likely blocked or restricted by coastal glaciers and geomorphic activity.

Thus, our earlier hypothesis (Khrustaleva et al., 2020) that the ancient Parakam Lake, existed in the area of the modern Kamakovo Lowland (Braitseva et al., 1968), and the network of its tributaries were one of the refugia in the Asian part of the sockeye salmon range, where a large population probably existed from the Sangamonian interglacial (∼ 130−115 ka) and throughout the last glaciation, can be considered convincing. In the sample from the Dvu’yurta River, the estimates of haplotype and nucleotide diversity were the highest. Probably, despite the fact that the river network underwent changes during the lake drying up and the river bed reorganization due to the melting of glaciers and the geological, tectonic and volcanic processes that occurred in the Holocene in this region, the river Dvu’yurta is a relict watercourse that once connected with the Parakam Lake, where the aboriginal population survived in the last glaciation has been preserved, and the early sockeye salmon of the Dvu’yurta River or, most likely, Dvu’yurta Lake at its source, can be confidently classified as glacial relics.

There is an understanding that the areas of the Gulf of Anadyr, the Anadyr River and some rivers of the Penzhina Lowland, were also not covered by ice sheet during the LGM, as evidenced by the diverse freshwater fauna in these water bodies (Kurenkov, 1965; Chereshnev, 1998; Altukhov et al., 2004). However, apparently, despite the fact that this region can be considered the largest refugium for the Arctic freshwater fish species in Northeast Asia (Chereshnev, 1998), we have no evidence that there was a conservation and dispersal area for sockeye salmon here during the last glaciation.

The regions south of the Kuril Islands also remained ice-free during the LGM. The first Illinois (Riss) glaciation (∼190–130 ka) covered all islands of the Greater Kuril Ridge and formed ice fields in the north and alpine glaciers in the south. The second (Wisconsin) glaciation left minimal traces on the central islands, with only small moraines observed above 1000–1200 m elevation starting from Urup Island. Similar small-scale Wisconsin moraines were identified on Iturup Island but are absent further south (Gorshkov, 1967). Moreover during the late Pleistocene regressions, the islands of Sakhalin, Hokkaido, Kunashir, the Lesser Kuril Islands and Iturup formed a common landmass connected to Primorye (Gorshkov, 1967; Chereshnev, 1998). We hypothesize that populations could have persisted in ice-free watersheds across this vast exposed shelf region – descendants of the first wave of colonization that migrated from northern territories and survived the Wisconsin glaciation. Previous studies have established that these same territories served as a major refugium and expansion center for chum salmon (*O. keta*) in Asia (Sato et al., 2004; Savin et al., 2009). The existence of a glacial refugium for sockeye salmon in the Hokkaido region is supported by the persistence of isolated relict populations of kokanee inhabiting lacustrine systems south of the contemporary range edge, as well as by the shared freshwater ichthyofauna and malacofauna between the Hokkaido and Iturup Islands. It is generally accepted that the ichthyofauna of the southern Kuril Islands represents a relict of a formerly more extensive southern refugium. Furthermore, biogeographic evidence suggests that the dispersal of modern fauna to the southern Kuril Islands from Hokkaido occurred during the Late Pleistocene, when land bridge connected these islands due to lower sea levels (Chereshnev, 1998; Sidorov & Pichugin, 2005; Bogatov et al., 2006). During the initial phase of the Holocene transgression, the temporary land bridge was flooded, prompting a northward range shift of sockeye salmon toward its ecological optimum in response to large-scale climatic warming. Some of the migrants could establish populations in suitable habitats in the southern Kuril Islands. Under this scenario the observed low haplotype diversity in Iturup Island samples is likely a result of a founder effect. Furthermore, the reduction in genetic diversity may be caused by prolonged isolation of relatively small island populations due to genetic drift or relatively recent bottleneck events. In particular, recent decline in population size was confirmed in Sopochnoye Lake according Tajima’s *D* tests. Our haplotype frequency distribution data suggest that the southern Kuril populations experienced minimal introgression from the northern populations in Holocene.

### Genealogy and Phylogeny of Haplotypes and Postglacial Expansion Patterns in Asia

Molecular dating enabled us to estimate the timing of substitutions and the divergence of the two Asian sockeye salmon lineages. The topology of the Bayesian tree suggests a relatively ancient origin of the central insertion/deletion (indel) at position 580 in the poly-T region of the D-loop. This indel is dated to approximately 290 thousand years ago, corresponding to the pre-Illinoian period or the Dnieper glaciation in the Northern Hemisphere (maximum ∼300–250 ka). The second indel, at position 845, is estimated to be younger, dating to roughly 110 ka. Around the same period (∼160 ka), during the Illinoian glacial stage (∼190–130 ka), the divergence of the sockeye salmon phylogenetic lineages occurred. Most modern haplotypes appear to have originated between 90–30 ka, coinciding with the Last Glacial Maximum (Wisconsin Glaciation, ∼75–11 ka).

Comparison of the three genealogies suggests that the key transitional haplotypes between the two phylogroups are Hap_11, Hap_23, and Hap_24. Their geographic distribution − found in Iturup Island and the Kamchatka River basin − may indicate the location of glacial refugia for Asian sockeye salmon. Additionally, in the southernmost part of the species’ Asian range, all three ancient haplotypes are present, with an estimated age of approximately 190–110 thousand years ago. This suggests that a relict "southern" population likely persisted in this region, near the glacial margin, during the Late Pleistocene regressions. This population may have been ancestral to contemporary sockeye salmon populations inhabiting the southern Kuril Islands and Hokkaido Island.

Estimating the emergence time of sockeye salmon populations in the South Kuril Islands and modeling postglacial expansion scenarios in this region would enable extrapolation of these findings to the broader western Pacific coast, thereby reconstructing the evolutionary history of the Asian lineage. Two alternative hypotheses may explain the origin of the Kuril populations (the third possible hypothesis about the colonization of the region during the pre-Illinoian interstadial (∼250–190 ka) is indefensible due to extreme difficulty of an appropriate scenario modeling and the lack of evidence for Asian populations existence in such a distant past):

1. Post-Illinoian Expansion hypothesis suggests that the Asian range expansion by North American /Beringian sockeye salmon populations occurred in the post-Illinoian time, during the Sangamonian interglacial. Bayesian divergence time estimates for the haplogroups (∼160-110 ka) coincide with this temporal window. The ancient transitional haplotypes likely originated on the American Pacific coast (∼190−160 ka), subsequently spreading throughout the Asian range during the transgression phase (∼130−80 ka), where they persisted in large refugial watersheds. During the LGM (∼75−11 ka), the 10T phylogenetic lineage emerged in the Kamchatka River basin through intraclade haplotype radiation. Secondary contact between previously isolated continental populations was then reestablished during the Holocene transgression.
2. Holocene Colonization hypothesis rejects the existence of a southern refugium, proposing instead that the South Kurils were colonized during the Holocene, contemporaneously with the southward expansion of Beringian and Kamchatka populations.

The proposed hypotheses (the first corresponding to scenarios 3 and 4, and the second to scenarios 1 and 2 showed in Fig. 6a) were evaluated through coalescent simulations and ABC−based assessment of historical inference probabilities implemented in DIYABC. ABC modeling results support a scenario of recurrent spatial expansions of sockeye salmon across the Asian range during the last two glacial cycles. Probably, the first wave of the species invasion to Asia, following the Middle Pleistocene Illinois glaciation (∼250−140 ka), led to the expansion of the American 13T haplotype along the entire Asian−Pacific coast up to the northern islands of the Japanese archipelago. However, subsequent Upper Pleistocene climatic oscillations again promoted range fragmentation, isolating populations in multiple refugia where they diverged through local adaptation and/or haplotype frequency shifts caused by genetic drift. The beginning of the Holocene transgression coincided with an extremely rapid (explosive) expansion of this species throughout its modern range. Furthermore watersheds were colonized contemporaneously by individuals from distinct geographic regions including northern (Alaska/Beringia) and southern (Hokkaido Island) populations synchronously with the radiation of the 10T haplotype from the central Kamchatka refugium. Analysis of SNP haplotype distribution along the North Pacific coast suggests that the 10T phylogenetic lineage invaded from the Kamchatka River basin to Cook Inlet and Alaska Peninsula river systems, likely using the Aleutian island chain as a migratory corridor (the "island bridge" hypothesis).

## CONCLUSION

Our findings support a primary scenario for the formation of contemporary mtDNA haplotype diversity in sockeye salmon in Asia implying multiple colonization events from North American (Beringian) populations during Pleistocene climatic optima. Post−LGM expansion sites on the Asian coast appear to have been located in the Kamchatka River basin refugium and potentially in a cryptic southern refugium in the Hokkaido region, though the genetic signature of this refugium was largely lost due to population fragmentation during glacial retreat. Biogeographic analysis of Kuril Islands colonization patterns indicates that both anadromous sockeye salmon populations in southern islands and kokanee populations in Sopochnoe Lake (Iturup Island) and Hokkaido watersheds represent glacial relicts that persisted through the LGM in southern refugia. Consequently, conserving the unique gene pools of these native island populations particularly under escalating anthropogenic pressure and climate change necessitates implementing targeted conservation strategies. These should incorporate comprehensive genetic monitoring programs and flexible fisheries management protocols.

## Acknowledgments

Author gratefully acknowledges to Dr. J.E. Seeb (College of the Environment, University of Washington, Seattle, WA, USA), as well as to all the employees of the Environmental Genomics Laboratory, Department of Hydrobiology and Fisheries, University of Washington, for their guidance and assistance in SNP genotyping, Klovach N.V., Dr.science (Russian Federal Research Institute of Fisheries and Oceanography –VNIRO), for sample collection management, Volkov A.A. (Syntol Research and Production Firm) for D−loop sequencing methods developing, primers design, and technical support, as well as the employees of A.N. Severtsov Institute of Ecology and Evolution of the Russian Academy of Sciences, VNIRO, Kamchatka branch (KamchatNIRO), Pacific branch (TINRO), Sakhalin branch (SakhNIRO), and Magadan branch (MagadanNIRO) of the VNIRO, who took part in the samples collection.

## Author Contributions

Anastasia M. Khrustaleva: conceptualisation; funding acquisition; supervision; methodology; investigation (equal); data curation; formal analysis; software; visualisation; writing – original draft preparation. Ekaterina V. Ponomareva: investigation (equal); writing – review. All authors read and approved the final version of the manuscript.

## Data Availability

The haplotype sequence data have been submitted to the GenBank database under accession IDs MT304834−MT304856 ("long-length") and PV892498-PV892517 ("short-length").

## Ethical approval

This study was carried out in compliance with the Federal Law No. 498−FZ ‘On Responsible Handling of Animals and on Amending Certain Legislative Acts of the Russian Federation’ (17 December 2018). No ethical approval was required for fish provided dead by local fisheries.

## Supplementary Information

The online version contains supplementary material available at …

## REFERENCES

Afanas’ev, K. I., G. A. Rubtsova, M. V. Shitova, T. V. Malinina, T. A. Rakitskaya, V. D. Prokhorovskaya, E. A. Shevlyakov, L. O. Zavarina, L. T. Bachevskaya, I. A. Chereshnev, V. A. Brykov, M. Y. Kovalev, V. A. Shevlyakov, S. V. Sidorova, S. I. Borzov, V. P. Pogodin, L. K. Fedorova, & L. A. Zhivotovsky, 2011. Population structure of chum salmon Oncorhynchus keta in the Russian Far East, as revealed by microsatellite markers. Russ. J. Mar. Biol. 37: 42–51.

Altukhov, Y. P., E. A. Salmenkova, & O. L. a al Kurbatova, 2004. Dinamika populyatsionnykh genofondov pri antropogennykh vozdeistviyakh (The dynamics of population gene pools under anthropogenic impacts). Nauka, Moscow.

Bachevskaya, L. T., G. D. Ivanova, V. V. Pereverzeva, & G. A. Agapova, 2017. Genetic structure of the coho salmon Oncorhynchus kisutch in the rivers of northeastern Russia according to the data on the variability of the cytochrome b gene of mitochondrial DNA. Biol. Bull. (Moscow) Maik Nauka Publishing / Springer SBM 44: 568–574.

Bachevskaya, L. T., V. V. Pereverzeva, & G. D. Ivanova, 2015. Genetic diversity of sockeye salmon (Oncorhynchus nerka) from some rivers of eastern Kamchatka and continental coast of the Sea of Okhotsk according to the data on cytochrome b gene polymorphism of mitochondrial DNA. Issled. Vodn. Biol. Resur. Kamchatki Sev.-Zapadn. Chasti Tikhogo Okeana 38: 49–56.

Bachevskaya, L. T., V. V. Pereverzeva, & T. V. Malinina, 2011. The genetic structure of chum salmon (Oncorhynchus keta Walbaum) populations inferred from the nucleotide variation of the mitochondrial DNA cytochrome b gene. Russ. J. Genet 47: 1314–1323.

Bachevskaya, L. T., V. V. Pereverzeva, A. A. Primak, & G. A. Agapova, 2021. Genetic Diversity of the Pink Salmon Oncorhynchus gorbuscha (Walbaum) from Some Rivers of Northeastern Russia. Biology Bulletin Pleiades journals 48: 413–424, https://link.springer.com/article/10.1134/S1062359021040038.

Beacham, T. D., B. McIntosh, C. MacConnachie, K. M. Miller, R. E. Withler, & N. Varnavskaya, 2006a. Pacific Rim population structure of sockeye salmon (Oncorhynchus nerka) as determined from microsatellite analysis. Transactions of the American Fisheries Society Wiley 135: 174–187.

Beacham, T. D., N. V. Varnavskaya, B. McIntosh, & C. MacConnachie, 2006b. Population Structure of Sockeye Salmon from Russia Determined with Microsatellite DNA Variation. Transactions of the American Fisheries Society Wiley 135: 97–109.

Becker, R. A., T. P. Minka, & A. Deckmyn, 2022. maps: Draw Geographical Maps., https://cran.r-project.org/package=maps.

Bermingham, E., S. S. Mccafferty, & A. P. Martin, 1997. Fish Biogeography and Molecular Clocks: Perspectives from the Panamanian Isthmus. Molecular Systematics of Fishes Elsevier 113–128.

Bickham, J. W., C. C. Wood, & J. C. Patton, 1995. biogeographic implications of cytochrome b sequences and allozymes in sockeye (Oncorhynchus nerka). Journal of Heredity Oxford University Press 86: 140–144.

Bogatov, V. V., T. W. Pietsch, S. Y. Storozhenko, V. Y. Barkalov, A. S. Leley, S. K. Kholin, P. V. Krestov, V. A. Kostenko, E. A. Makarchenko, L. A. Prozorova, & S. V. Shedko, 2006. Origin patterns of the terrestrial and freshwater biota of Sakhalin Island. Vestnik DVO (Bulletin of the Russian Academy of Sciences, Far East Branch) 2: 32–47.

Bouckaert, R., T. G. Vaughan, J. Barido-Sottani, S. Duchêne, M. Fourment, A. Gavryushkina, J. Heled, G. Jones, D. Kühnert, N. De Maio, M. Matschiner, F. K. Mendes, N. F. Müller, H. A. Ogilvie, L. Du Plessis, A. Popinga, A. Rambaut, D. Rasmussen, I. Siveroni, M. A. Suchard, C. H. Wu, D. Xie, C. Zhang, T. Stadler, & A. J. Drummond, 2019. BEAST 2.5: An advanced software platform for Bayesian evolutionary analysis. PLOS Computational Biology Public Library of Science 15: e1006650, https://journals.plos.org/ploscompbiol/article?id=10.1371/journal.pcbi.1006650.

Braitseva, O. A., & I. S. Evteeva, 1968. Climatic fluctuations and the Pleistocene glaciations of Kamchatka. Geol. Geofiz. 5: 16–22.

Braitseva, O. A., I. V. Melekestsev, I. S. Evteeva, & E. G. Lupikina, 1968. Stratigrafiya chetvertichnykh otlozhenii i oledeneniya Kamchatki (Stratigraphy of the Quaternary sediments and glaciations of Kamchatka). Nauka, Moscow.

Brunner, P. C., M. R. Douglas, A. Osinov, C. C. Wilson, & L. Bernatchez, 2001. Holarctic phylogeography of arctic charr (Salvelinus alpinus L.) Inferred from mitochondrial DNA sequences. Evolution Society for the Study of Evolution 55: 573–586.

Brykov, V. A., N. E. Polyakova, A. V. Podlesnykh, E. V. Golub’, A. P. Golub’, & O. L. Zhdanova, 2005. The effect of reproduction biotopes on the genetic differentiation of populations of sockeye salmon Oncorhynchus nerka. Russ. J. Genet. 41: 509–517.

Bugaev, V. F., & V. E. Kirichenko, 2008. Nagulno-nerestovye ozera aziatskoi nerki (vklyuchaya nekotorye drugie vodoemy areala) (The feeding-spawning lakes of the Asian sockeye salmon (including some other reservoirs of the range). Kamchatpress, Petropavlovsk-Kamchatsky.

CCU“Genome,” 2011. Sample preparation., http://www.genome-centre.ru/preparation.html.

Chereshnev, I. A., 1998. Biogeografiya presnovodnykh ryb Dal’nego Vostoka Rossii. Dal’nauka, Vladivostok, Russia.

Christensen, K. A., E. B. Rondeau, D. R. Minkley, D. Sakhrani, C. A. Biagi, A. M. Flores, R. E. Withler, S. A. Pavey, T. D. Beacham, T. Godin, E. B. Taylor, M. A. Russello, R. H. Devlin, & B. F. Koop, 2020. The sockeye salmon genome, transcriptome, and analyses identifying population defining regions of the genome. PLoS ONE., 10.1371/journal.pone.0240935.

Churikov, D., M. Matsuoka, X. Luan, A. K. Gray, V. L. A. Brykov, & A. J. Gharrett, 2001. Assessment of concordance among genealogical reconstructions from various mtDNA segments in three species of Pacific salmon (genus Oncorhynchus). Mol. Ecol. 10: 2329–2339.

Cline, T. J., J. Ohlberger, & D. E. Schindler, 2019. Effects of warming climate and competition in the ocean for life-histories of Pacific salmon. Nature Ecology and Evolution Nature Publishing Group 3: 935–942, https://www.nature.com/articles/s41559-019-0901-7.

Collin, F. D., G. Durif, L. Raynal, E. Lombaert, M. Gautier, R. Vitalis, J. M. Marin, & A. Estoup, 2021. Extending approximate Bayesian computation with supervised machine learning to infer demographic history from genetic polymorphisms using DIYABC Random Forest. Molecular Ecology Resources John Wiley & Sons, Ltd 21: 2598–2613, https://onlinelibrary.wiley.com/doi/full/10.1111/1755-0998.13413.

Crozier, L. G., B. J. Burke, B. E. Chasco, D. L. Widener, & R. W. Zabel, 2021. Climate change threatens Chinook salmon throughout their life cycle. Communications Biology Nature Research 4:.

Dellicour, S., & P. Mardulyn, 2014. spads 1.0: a toolbox to perform spatial analyses on DNA sequence data sets. Molecular Ecology Resources John Wiley & Sons, Ltd 14: 647–651, https://onlinelibrary.wiley.com/doi/full/10.1111/1755-0998.12200.

Drummond, A. J., A. Rambaut, B. Shapiro, & O. G. Pybus, 2005. Bayesian coalescent inference of past population dynamics from molecular sequences. Molecular Biology and Evolution 22: 1185–1192.

Excoffier, L., & H. E. L. Lischer, 2010. Arlequin suite ver 3.5: A new series of programs to perform population genetics analyses under Linux and Windows. Molecular Ecology Resources 10: 564–567.

Gearty, W., 2024. deeptime: an R package that facilitates highly customizable visualizations of data over geological time intervals. EarthArXiv.

Gorshkov, G. S., 1967. Vulkanizm Kuril’skoi ostrovnoi dugi. Moscow: Nauka, Moscow: Nauka.

Grant, W. S., 2015. Problems and cautions with sequence mismatch analysis and Bayesian skyline plots to infer historical demography. Journal of Heredity Oxford University Press 106: 333–346.

Habicht, C., L. W. Seeb, K. W. Myers, E. V. Farley, & J. E. Seeb, 2010. Summer–Fall Distribution of Stocks of Immature Sockeye Salmon in the Bering Sea as Revealed by Single-Nucleotide Polymorphisms. Transactions of the American Fisheries Society Wiley 139: 1171–1191.

Hasegawa, M., H. Kishino, & T. aki Yano, 1985. Dating of the human-ape splitting by a molecular clock of mitochondrial DNA. Journal of molecular evolution J Mol Evol 22: 160–174, https://pubmed.ncbi.nlm.nih.gov/3934395/.

Hewitt, G. M., 1996. Some genetic consequences of ice ages, and their role in divergence and speciation. Biological Journal of the Linnean Society Oxford University Press (OUP) 58: 247–276.

Jenkins, T. L., R. Castilho, & J. R. Stevens, 2018. Meta-analysis of northeast Atlantic marine taxa shows contrasting phylogeographic patterns following post-LGM expansions. PeerJ PeerJ Inc. 2018:.

Khrustaleva, A. M., 2016. The phylogeography of the asian sockeye salmon Oncorhynchus nerka, inferred from the data on the variability of mitochondrial SNP loci: Analysis of scenarios for post-glacial expansion of the species over the asian coast of the pacific ocean. Russian Journal of Marine Biology Maik Nauka Publishing / Springer SBM 42: 517–526, https://link.springer.com/article/10.1134/S1063074016070051.

Khrustaleva, A. M., E. V. Ponomareva, M. V. Ponomareva, E. A. Shubina, & O. A. Pilganchuk, 2020. Phylogeography of Asian sockeye salmon (Oncorhynchus nerka) based on analysis of mtDNA control region polymorphism. Journal of Applied Ichthyology 36: 643–654.

Kondzela, C., & A. J. Gharrett, 2007. Preliminary analysis of sockeye salmon colonization in Glacier Bay inferred from genetic methods. In Piatt, J. F., & S. M. Gende (eds), Proceedings of the Fourth Glacier Bay Science Symposium, October 26–28, 2004: U. S. Geological Survey Scientific Investigations Report 2007-5047. : 110–114.

Koval, M. V. V., O. B. B. Tepnin, S. L. L. Gorin, E. S. S. Fadeev, E. V. Zikunova, O.V. Lepskaya, S. B. Shubkin, S.V. Rudakova, S.L. Pilganchuk, O.A. Gorodovskaya, O. V. Zikunova, E. V. Lepskaya, S. V. Shubkin, S. L. Rudakova, O. A. Pilganchuk, & S. B. Gorodovskaya, 2020. Factors determining spawning run dynamics and current state of sockeye salmon Oncorhynchus nerka resources in the Kamchatka River. The researches of the aquatic biological resources of Kamchatka and the North-West Part of the Pacific Ocean 57: 5–66.

Kurenkov, I. I., 1965. Zoogeography of Freshwater Fish from Kamchatka. Vopr. Geogr. Kamchatki 3: 25–34.

Marchenko, S. L., 2022. Sockeye salmon Oncorhynchus nerka (Salmoniformes, Salmonidae) of continental coast of the Sea of Okhotsk. Problems of Fisheries 23(3): 102–121.

McCusker, M. R., E. Parkinson, & E. B. Taylor, 2000. Mitochondrial DNA variation in rainbow trout (Oncorhynchus mykiss) across its native range: testing biogeographical hypotheses and their relevance to conservation. Molecular Ecology John Wiley & Sons, Ltd 9: 2089–2108, https://onlinelibrary.wiley.com/doi/full/10.1046/j.1365-294X.2000.01121.x.

Nguyen, L. T., H. A. Schmidt, A. Von Haeseler, & B. Q. Minh, 2015. IQ-TREE: A Fast and Effective Stochastic Algorithm for Estimating Maximum-Likelihood Phylogenies. Molecular Biology and Evolution Oxford Academic 32: 268–274, 10.1093/molbev/msu300.

Nikulin, O. A., 1975. Reproduction of red Oncorhynchus nerka (Walb.) in the Okhota River basin. Tr. Vseross. Nauchno-Issled. Inst. Rybn. Khoz. Okeanogr. CVI: 97–105.

Oke, K. B., C. J. Cunningham, P. A. H. Westley, M. L. Baskett, S. M. Carlson, J. Clark, A. P. Hendry, V. A. Karatayev, N. W. Kendall, J. Kibele, H. K. Kindsvater, K. M. Kobayashi, B. Lewis, S. Munch, J. D. Reynolds, G. K. Vick, & E. P. Palkovacs, 2020. Recent declines in salmon body size impact ecosystems and fisheries. Nature Communications Nature Research 11: 1–13, https://www.nature.com/articles/s41467-020-17726-z.

Paradis, E., & J. Barrett, 2010. pegas: an R package for population genetics with an integrated–modular approach. Bioinformatics Oxford Academic 26: 419–420, 10.1093/bioinformatics/btp696.

Pebesma, E., & R. Bivand, 2023. Spatial Data Science: With Applications in R. Chapman and Hall/CRC, https://r-spatial.org/book/.

Prozorova, L. A., & M. B. Shedko, 2003. Mollusks of Lake Azabachye and their biocenotic significance. Transactions of the Kamchatka Branch of the Pacific Institute of Geography, Far Eastern Branch of the Russian Academy of Sciences. 4: 120–151.

Rambaut, A., 2018. Figtree ver 1.4.4. Institute of Evolutionary Biology, University of Edinburgh. Edinburgh.

Rambaut, A., A. J. Drummond, D. Xie, G. Baele, & M. A. Suchard, 2018. Posterior Summarization in Bayesian Phylogenetics Using Tracer 1.7. Systematic Biology Oxford Academic 67: 901–904, 10.1093/sysbio/syy032.

Ramos-Onsins, S. E., & J. Rozas, 2002. Statistical Properties of New Neutrality Tests Against Population Growth. Molecular Biology and Evolution Oxford Academic 19: 2092–2100, 10.1093/oxfordjournals.molbev.a004034.

Rogers, A. R., & H. Harpending, 1992. Population growth makes waves in the distribution of pairwise genetic differences. Molecular Biology and Evolution Society for Molecular Biology and Evolution 9: 552–569.

Sato, S., H. Kojima, J. Ando, H. Ando, R. L. Wilmot, L. W. Seeb, V. Efremov, L. LeClair, W. Buchholz, D. H. Jin, S. Urawa, M. Kaeriyama, A. Urano, & S. Abe, 2004. Genetic population structure of chum salmon in the Pacific Rim inferred from mitochondrial DNA sequence variation. Environmental Biology of Fishes 69: 37–50.

Savin, V. A., N. Y. Shpigalskaya, & N. V. Varnavskaya, 2009. Interregional and interpopulation variations of chum salmon (Oncorhynchus keta) mitochondrial DNA haplotype frequencies in Asia. The researches of the aquatic biological resources of Kamchatka and of the northwest part of the Pacific Ocean 12: 15–32.

Seeb, J. E., C. E. Pascal, R. Ramakrishnan, & L. W. Seeb, 2009. SNP genotyping by the 5’-nuclease reaction: advances in high-throughput genotyping with nonmodel organisms. Methods in molecular biology (Clifton, N.J.) 578: 277–292.

Shafer, A. B. A., C. I. Cullingham, S. D. Côté, & D. W. Coltman, 2010. Of glaciers and refugia: a decade of study sheds new light on the phylogeography of northwestern North America. Molecular Ecology John Wiley & Sons, Ltd 19: 4589–4621, /doi/pdf/10.1111/j.1365-294X.2010.04828.x.

Shpigalskaya, N. Y., V. A. Brykov, A. J. Gharrett, A. D. Kukhlevsky, R. A. Shaporev, & N. V. Varnavskaya, 2008. Variation of mitochondrial DNA in chinook salmon Oncorhynchus tschawytscha Walbaum populations from Kamchatka. Russian Journal of Genetics Springer 44: 849–858, https://link.springer.com/article/10.1134/S1022795408070132.

Shpigalskaya, N. Y., V. A. Brykov, & A. D. Kukhlevsky, 2009. Pink salmon mtDNA polymorphysm in Kamchatka and Sakhalin. The researches of the aquatic biological resources of Kamchatka and of the northwest part of the Pacific Ocean 13: 74–87.

Sidorov, L. K., & M. Y. Pichugin, 2005. Composition of the ichthyofauna and specifics of the biology of fish from the Southern Kuril Islands in connection with abiotic conditions and the origin of water bodies. Trudy VNIRO 144: 151–175.

Smith, G. R., 1992. Introgression in Fishes: Significance for Paleontology, Cladistics, and Evolutionary Rates. Systematic Biology Oxford Academic 41: 41–57, 10.1093/sysbio/41.1.41.

Stepien, C. A., A. K. Dillon, & M. J. Brooks, 1997. The evolution of Blennioid fishes based on analysis of mitochondrial 12S rDNA., 245–270.

Thompson, T. Q., M. Renee Bellinger, S. M. O’Rourke, D. J. Prince, A. E. Stevenson, A. T. Rodrigues, M. R. Sloat, C. F. Speller, D. Y. Yang, V. L. Butler, M. A. Banks, & M. R. Miller, 2019. Anthropogenic habitat alteration leads to rapid loss of adaptive variation and restoration potential in wild salmon populations. Proceedings of the National Academy of Sciences of the United States of America National Academy of Sciences 116: 177–186, /doi/pdf/10.1073/pnas.1811559115?download=true.

Varnavskaya, N. V., 2006. Geneticheskaya differentsiatsiya populyatsii tikhookeanskikh lososei. Publishing house KamchatNIRO, Petropavlovsk-Kamchatsky.

Varnavskaya, N. V., C. C. Wood, & R. J. Everett, 1994. Genetic variation in sockeye salmon (Oncorhynchus nerka) populations of Asia and North America. Can. J. Fish. Aquat. Sci Canadian Science Publishing 51: 132–146.

Velichko, A. A., & M. A. Faustova, 1989. Rekonstruktsii poslednego pozdnepleistotsenovogo oledeneniya Severnogo polushariya (18–20 tys. let nazad). (Reconstruction of the last Late Pleistocene glaciation of the Northern Hemisphere (18–20 thousand years ago). Doklady AN SSSR 309: 1465–1468.

Villesen, P., 2007. FaBox: an online toolbox for fasta sequences. Molecular Ecology Notes John Wiley & Sons, Ltd 7: 965–968, https://onlinelibrary.wiley.com/doi/full/10.1111/j.1471-8286.2007.01821.x.

Waples, R. S., R. W. Zabel, M. D. Scheuerell, & B. L. Sanderson, 2008. Evolutionary responses by native species to major anthropogenic changes to their ecosystems: Pacific salmon in the Columbia River hydropower system. Molecular Ecology 17: 84–96.

Waterhouse, A. M., J. B. Procter, D. M. A. Martin, M. Clamp, & G. J. Barton, 2009. Jalview Version 2—a multiple sequence alignment editor and analysis workbench. Bioinformatics Oxford Academic 25: 1189–1191, 10.1093/bioinformatics/btp033.

Wickham, H., 2009. ggplot2 Elegant Graphics for Data Analysis. ggplot2. Springer New York, New York, NY, http://link.springer.com/10.1007/978-0-387-98141-3.

Wood, C. C., B. E. Riddell, D. T. Rutherford, & R. E. Withler, 1994. Biochemical Genetic Survey of Sockeye Salmon (Oncorhynchus nerka) in Canada. Canadian Journal of Fisheries and Aquatic Sciences 51: 114–131.

Xu, S., L. Li, X. Luo, M. Chen, W. Tang, L. Zhan, Z. Dai, T. T. Lam, Y. Guan, & G. Yu, 2022. Ggtree: A serialized data object for visualization of a phylogenetic tree and annotation data. iMeta John Wiley & Sons, Ltd 1: e56, https://onlinelibrary.wiley.com/doi/full/10.1002/imt2.56.

Zelenina, D. A., V. A. Soshnina, & A. A. Sergeev, 2020. Phylogeography and Mitochondrial Polymorphism of Asian Coho Salmon. Molekuliarnaia biologiia 54: 997–1005.

Zhivotovsky, L. A., S. D. Pavlov, M. Y. Kovalev, V. A. Parensky, E. V. Ponomareva, M. N. Mel’nikova, T. V. Mineeva, A. L. Senchukova, T. A. Rakitskaya, G. A. Rubtsova, & K. I. Afanasyev, 2019. Genetic Differentiation of the Resident and Anadromous Sockeye Salmon Populations of the Kamchatka Peninsula: An Evolutionary Scenario for the Origin of the Resident Sockeye Salmon in Lake Kronotskoye. Russian Journal of Marine Biology Pleiades Publishing 45: 443–452.

