## Supplementary material for "Linking Ancient Refugia to Modern Diversity: Evidence of Multi-origin Postglacial Expansion of Sockeye Salmon on the Asian Range": Table S1

**Table S1.** Prior settings of four biogeographical scenarios, which were tested under an ABC framework

| Sce−nario | Prior settings of ranges for time intervals | | | Prior settings of effective population size (*Ne*) | | |
|---|---|---|---|---|---|---|
| | Designa−tions | Descriptions | Time, years | Designations | Descriptions | *Ne*, number of fish |
| 1 | $t_1$ − $t_4$ | Time to population divergence | 1 − 26,500; $t_1 \le t_2 \le t_3 \le t_4$ | N1 | Contemporary Kuril Lake population *Ne* | 100,000 – 1,500,000 |
| | db ($t_{db}$) | Time to population size change | 10 − 26,500; $t_{db} \le t_1 \ldots t_4$ | N2 – N5 | Contemporary Kuril Islands populations' *Ne* | 500 – 5,000 |
| | | | | $N2_b$ − $N5_b$ | Invasive population size | 10 – 1,000 |
| 2 | $t_1$ − $t_4$ | Same as Scenario1 | 1 − 26,500; $t_1 \le t_2 \le t_3 \le t_4$ | N1 | Same as Scenario 1 | 100,000 – 1,500,000 |
| | db ($t_{db}$) | Same as Scenario1 | 10 − 26,500; $t_{db} \le t_1 \ldots t_4$ | N2 – N5 | Same as Scenario 1 | 500 – 5,000 |
| | | | | $N2_b$ − $N5_b$ | Same as Scenario 1 | 10 – 1,000 |
| | | | | $N1_c$ | Ancestral invasive population size | 100 – 10,000 |
| 3 | $t_1$ – $t_3$ | Same as Scenario1 | 1 − 26,500; $t_1 \le t_2 \le t_3 \le t_4$ | N1 | Same as Scenario 1 | 100,000 – 1,500,000 |
| | db ($t_{db}$) | Same as Scenario1 | 10 − 26,500; $t_{db} \le t_1 \ldots t_4$ | N2 – N5 | Same as Scenario 1 | 500 – 5,000 |
| | $t_5$ | Time to post−Illinoian expansion | 135,000 – 190,000 | $N2_b$ − $N5_b$ | Same as Scenario 1 | 10 – 1,000 |
| | | | | $N1_c$ | Same as Scenario 2 | 100 – 10,000 |
| | | | | Na | Ancestral population size | 100,000 – 1,500,000 |
| 4 | $t_1$ − $t_4$ | Same as Scenario1 | 1 − 26,500; $t_1 \le t_2 \le t_3 \le t_4$ | N1 | Same as Scenario 1 | 100,000 – 1,500,000 |
| | db ($t_{db}$) | Same as Scenario1 | 10 − 26,500; $t_{db} \le t_1 \ldots t_4$ | N2 – N5 | Same as Scenario 1 | 500 – 5,000 |
| | $t_5$ | Same as Scenario 3 | 135,000 – 190,000 | $N2_b$ − $N5_b$ | Same as Scenario 1 | 10 – 1,000 |
| | $t_a$ | Time to admixture event | 10 − 26,500; $t_1 \le t_2 \le t_3 \le t_4 \le t_a$ | $N1_c$ | Same as Scenario 2 | 100 – 10,000 |
| | | | | $N1_b$ | Ancestral population size in Holocene | 100,000 – 1,500,000 |
| | | | | Na | Ancestral population size in post−Illinois | 100,000 – 1,500,000 |

**Table S2**. Samples characteristics, regions and locations, population IDs, date of catch, etc. and mtSNP haplotype frequencies of Asian sockeye salmon populations

| # | Region | Location | Data source | Pop ID | *n* | Date | Coordi-nates | *One_CO1/One_Cytb_17/One_Cytb_26* (mtDNA haplotypes) | | | | | | |
|---|---|---|---|---|---|---|---|---|---|---|---|---|---|---|
| | | | | | | | | CGG | TGA | CAG | TAA | TGG | CGA | TAG |
| 1 | Chukotka, Navarinsky region | Vaamochka Lake | our data | Ch | 50 | 28.07.2004 | 62.54, 176.813 | 44.68 | 53.19 | 0 | 2.13 | 0 | 0 | 0 |
| 2 | Kamchatka peninsula, Olyutorsky region | Severnaya Lagoon | Habicht et al. 2010 | KSL | 98 | 26.06.2002 | 60.491, 170.812 | 73 | 27 | 0 | 0 | 0 | 0 | 0 |
| 3 | | Anana Lagoon | Habicht et al. 2010 | KAL | 80 | 24.06.2002 | 60.071, 170.241 | 65.4 | 33.3 | 0 | 0 | 0 | 1.3 | 0 |
| 4 | | Apuka River, early run | our data | KAerl | 18 | 24.06.2008-25.06.2008 | 60.486, 170.359 | 28.13 | 71.88 | 0 | 0 | 0 | 0 | 0 |
| 5 | | Apuka River, late run | our data | KAlt | 28 | 24.06.2008-25.06.2008 | 60.451, 169.673 | 53 | 47 | 0 | 0 | 0 | 0 | 0 |
| 6 | | Apuka River, Lake Vatit | Habicht et al. 2010 | KAvat | 51 | 07.08.2002 | 60.486, 171.359 | 22 | 78 | 0 | 0 | 0 | 0 | 0 |
| 7 | | Pakhacha River | our data | KPh | 59 | 17.06.2005-27.06.2005 | 60.907, 169.071 | 40.68 | 59.32 | 0 | 0 | 0 | 0 | 0 |
| 8 | | Pakhacha River, Lake Potat | Habicht et al. 2010 | KPhpot | 50 | 29.07.2001 | 60.681, 167.61 | 48 | 52 | 0 | 0 | 0 | 0 | 0 |
| 9 | Kamchatka peninsula, Karaginsky region | Khaylulya River | Bachevskaya et al. 2015 | KHay | 48 | no data | 58.183, 161.936 | 58 | 42 | 0 | 0 | 0 | 0 | 0 |
| 10 | Kamchatka River basin | Kamchatka River, late run | our data | KK-04 | 82 | 29.06.2004-09.07.2004 | 56.252, 162.42 | 80.85 | 19.15 | 0 | 0 | 0 | 0 | 0 |
| 11 | | Kamchatka River, early run | our data | KK-05 | 15 | 14.06.2005 | 56.252, 163.42 | 66.67 | 33.33 | 0 | 0 | 0 | 0 | 0 |
| 12 | | Kamchatka River, late run | Habicht et al. 2010 | KKerl-98 | 78 | 01.06.1998 | 56.236, 162.142 | 79 | 21 | 0 | 0 | 0 | 0 | 0 |
| 13 | | Kamchatka River, early run | Habicht et al. 2010 | KKlt-98 | 100 | 21.07.1998 | 56.236, 163.142 | 88 | 8 | 0 | 0 | 1 | 3 | 0 |
| 14 | | Azabachje Lake | our data | KKa | 81 | 03.07.2004, 13.07.2004 | 56.147, 161.8 | 53.25 | 46.75 | 0 | 0 | 0 | 0 | 0 |
| 15 | | Hapiza River | Habicht et al. 2010 | KKhap | 146 | 02.09.1998 | 56.042, 161.227 | 58 | 42 | 0 | 0 | 0 | 0 | 0 |
| 16 | | Dvu'yurta River | Habicht et al. 2010 | KKdv | 88 | 1994, 1995 | 56.81, 160.406 | 84 | 16 | 0 | 0 | 0 | 0 | 0 |
| 17 | | Elovka River | Habicht et al. 2010 | KKel | 109 | 1994, 1995 | 56.767, 160.827 | 90 | 10 | 0 | 0 | 0 | 0 | 0 |
| 18 | | Belaya River | Habicht et al. 2010 | KKbel | 88 | 1994, 1995 | 56.548, 160.145 | 88 | 12 | 0 | 0 | 0 | 0 | 0 |
| 19 | | Kozyrevka River | Habicht et al. 2010 | KKkoz | 40 | 1994 | 55.693, 159.433 | 79 | 21 | 0 | 0 | 0 | 0 | 0 |
| 20 | | Kitilgina River | Habicht et al. 2010 | KKkit | 28 | 29.06.1998 | 54.987, 159.121 | 54 | 46 | 0 | 0 | 0 | 0 | 0 |
| 21 | Kamchatka peninsula, East coast | Avacha River | Habicht et al. 2010 | KAv | 60 | 2002 | 53.447, 158.173 | 34 | 66 | 0 | 0 | 0 | 0 | 0 |

| | | | | | | | | | | | | | | |
|---|---|---|---|---|---|---|---|---|---|---|---|---|---|---|
| 22 | Commander Islands | Bering Island, Sarannoye Lake | our data | BS | 58 | 7.2008 | 55.276, 166.137 | 100 | 0 | 0 | 0 | 0 | 0 | 0 |
| 23 | Continental coast of the Sea of Okhotsk | Ola River | Bachevskaya et al. 2015 | Ola | 48 | no data | 59.659, 151.301 | 45 | 55 | 0 | 0 | 0 | 0 | 0 |
| 24 | | Okhota River | our data | Okh | 80 | 22.07.2004 | 59.469, 142.953 | 78.48 | 21.52 | 0 | 0 | 0 | 0 | 0 |
| 25 | Kamchatka peninsula, North-West | Palana River | Habicht et al. 2010 | KP-02 | 50 | 27.06.2002 | 59.078, 159.862 | 50 | 48.91 | 0 | 0 | 1.087 | 0 | 0 |
| 26 | | Palana River | our data | KP-03 | 94 | 10.07.2003-21.07.2003 | 59.078, 160.862 | 53.01 | 46.99 | 0 | 0 | 0 | 0 | 0 |
| 27 | | Tigil River | Habicht et al. 2010 | KT | 107 | 18.06.2002 | 57.981, 158.313 | 55 | 45 | 0 | 0 | 0 | 0 | 0 |
| 28 | Kamchatka peninsula, West coast | Vorovskaya River | our data | KV | 45 | 17.07.2007-27.07.2007 | 54.251, 155.819 | 62.75 | 37.25 | 0 | 0 | 0 | 0 | 0 |
| 29 | | Bolshaya River | our data | KB-03 | 91 | 23.07.2003-30.07.2003 | 52.628, 156.262 | 61.36 | 38.64 | 0 | 0 | 0 | 0 | 0 |
| 30 | | Bolshaya River | our data | KB-04 | 90 | 11.08.2004-20.08.2004 | 52.718, 157.224 | 62.07 | 36.78 | 1.15 | 0 | 0 | 0 | 0 |
| 31 | | Bolshaya River drainage, Bistraya River | our data | KBb-04 | 33 | 20.07.2004-12.08.2004 | 52.999, 157.747 | 72.73 | 27.27 | 0 | 0 | 0 | 0 | 0 |
| 32 | | Bolshaya River drainage, Bistraya River | Habicht et al. 2010 | KBb-98 | 56 | 16.08.1998 | 52.999, 156.747 | 67 | 29 | 0 | 0 | 2 | 2 | 0 |
| 33 | | Bolshaya River drainage, Plotnikova River | our data | KBp | 39 | 09.08.2004-12.082004 | 53.1, 157.757 | 60.53 | 39.47 | 0 | 0 | 0 | 0 | 0 |
| 34 | | Opala River | our data | KOp-07 | 50 | 01.07.2007 | 52.132, 156.477 | 48.98 | 51.02 | 0 | 0 | 0 | 0 | 0 |
| 35 | | Opala River | our data | KOp-08 | 31 | 17.07.2008-26.08.2008 | 52.132, 157.477 | 51.72 | 48.28 | 0 | 0 | 0 | 0 | 0 |
| 36 | | Ozernaya River | our data | KO | 95 | 04.08.2003-07.08.2003 | 51.501, 156.499 | 46.24 | 52.69 | 1.08 | 0 | 0 | 0 | 0 |
| 37 | North Kuril Islands | Shumshu Island, Bettobu Lake | our data | NKS | 50 | 05.08.2008 | 50.751, 156.264 | 48.94 | 51.06 | 0 | 0 | 0 | 0 | 0 |
| 38 | | Paramushir Island, Glukhoye Lake | our data | NKP | 48 | 07.07.2008-13.07.2008 | 50.489, 155.847 | 6.25 | 93.75 | 0 | 0 | 0 | 0 | 0 |
| 39 | South Kuril Islands | Urup Island, Tokotan Lake | our data | SKU | 35 | 07.2008-08.2008 | 45.859, 149.799 | 2.78 | 97.22 | 0 | 0 | 0 | 0 | 0 |
| 40 | | Iturup Island, Krasivoye Lake | our data | SKI | 50 | 01.10.2006 | 44.625, 147.209 | 14 | 86 | 0 | 0 | 0 | 0 | 0 |

**Table S3**. Samples characteristics, regions and locations, population IDs, date of catch, etc. and mtSNP haplotype frequencies of North-American sockeye salmon stocks

| # | Region | Collection name | Col. No | Date, m/dd/yyy | Coordinates | *n* | *One_CO1/One_Cytb_17/One_Cytb_26* (mtDNA haplotypes) | | | | | | |
|---|---|---|---|---|---|---|---|---|---|---|---|---|---|
| | | | | | | | CGG | TGA | CAG | TAA | TGG | CGA | TAG |
| 1 | Norton Sound | Glacial Lake | 27 | 8/15/2004 | 64.838, -165.705 | 96 | 0.189 | 0.789 | 0 | 0 | 0.021 | 0 | 0 |
| 2 | | Salmon Lake | 28 | 8/03/2001 | 64.896, -165.08 | 96 | 0.125 | 0.875 | 0 | 0 | 0 | 0 | 0 |
| 3 | Yukon River | Andreafsky River weir | 29 | 7/12/2005 | 62.118, -162.806 | 95 | 0.196 | 0.804 | 0 | 0 | 0 | 0 | 0 |
| 4 | | Gisasa River weir | 30 | 7/16/2005 | 65.258, -157.684 | 65 | 0.266 | 0.734 | 0 | 0 | 0 | 0 | 0 |
| 5 | Kuskokwim Bay and River | Kanektok River weir | 32 | 7/16/2002 | 59.75, -161.926 | 95 | 0.234 | 0.766 | 0 | 0 | 0 | 0 | 0 |
| 6 | | Salmon River - Aniak Basin | 35 | 8/02/2006 | 61.063, -159.198 | 95 | 0.211 | 0.789 | 0 | 0 | 0 | 0 | 0 |
| 7 | | Upper Takotna River | 37 | 2006 | 62.97, -155.613 | 40 | 0.216 | 0.784 | 0 | 0 | 0 | 0 | 0 |
| 8 | Togiak Drainage | Kulukak River Lake | 39 | 8/24/2006 | 59.171, -159.686 | 95 | 0.191 | 0.809 | 0 | 0 | 0 | 0 | 0 |
| 9 | Wood-Igushik Drainage | A Beach - Little Togiak Lake | 43 | 8/10/2005 | 59.567, -159.117 | 95 | 0.118 | 0.882 | 0 | 0 | 0 | 0 | 0 |
| 10 | Nushagak River | King Salmon River | 59 | 8/18/2001 | 60.251, -157.363 | 382 | 0.184 | 0.797 | 0 | 0 | 0.014 | 0.005 | 0 |
| 11 | Six Mile/Lake Clark | Chulitna Lodge Beach | 66 | 10/05/1999 | 60.181, -154.565 | 100 | 0.315 | 0.5 | 0 | 0 | 0.098 | 0.087 | 0 |
| 12 | Kvichak-Iliamna Drainage | Chinkelyes Creek | 71 | 8/28/2000 | 59.749, -153.87 | 275 | 0.344 | 0.652 | 0 | 0 | 0.004 | 0 | 0 |
| 13 | Naknek Drainage | Brooks Lake | 86 | 8/22/2000 | 58.493, -155.863 | 230 | 0.32 | 0.68 | 0 | 0 | 0 | 0 | 0 |
| 14 | Egegik Drainage | Becharof Creek | 93 | 8/11/2000 | 57.781, -155.955 | 660 | 0.433 | 0.562 | 0 | 0 | 0.002 | 0 | 0.003 |
| 15 | Ugashik Drainage | Black Creek | 96 | 8/24/2005 | 57.453, -156.715 | 382 | 0.478 | 0.516 | 0 | 0 | 0 | 0 | 0.005 |
| 16 | Peninsula - North | L Creek - Meshik River | 104 | 7/30/2005 | 56.749, -157.97 | 188 | 0.657 | 0.289 | 0 | 0 | 0 | 0 | 0.054 |
| 17 | | Ilnik River | 108 | 7/29/2002 | 56.585, -159.623 | 191 | 0.613 | 0.387 | 0 | 0 | 0 | 0 | 0 |
| 18 | | Davids River | 116 | 7/31/2005 | 55.764, -161.749 | 95 | 0.889 | 0.111 | 0 | 0 | 0 | 0 | 0 |
| 19 | Aleutian Islands | McLees Lake - Dutch Harbor | 119 | 6/04/2004 | 53.985, -166.726 | 95 | 0.67 | 0.33 | 0 | 0 | 0 | 0 | 0 |
| 20 | Peninsula - South | Thin Point Lagoon - near Cold Bay | 130 | 8/01/2005 | 55.002, -162.392 | 95 | 0.84 | 0.16 | 0 | 0 | 0 | 0 | 0 |
| 21 | Chignik | Chignik River | 137 | 8/22/1998 | 56.272, -158.663 | 95 | 0.457 | 0.543 | 0 | 0 | 0 | 0 | 0 |
| 22 | Kodiak/Afognak Islands | Afognak Lake | 144 | 8/15/1993 | 58.133, -152.986 | 80 | 0.474 | 0.423 | 0 | 0 | 0 | 0 | 0.103 |
| 23 | | Pasagshak Lake | 156 | 7/15/2005 | 57.473, -152.466 | 95 | 0.474 | 0.189 | 0 | 0 | 0 | 0 | 0.337 |
| 24 | Cook Inlet - West | Beluga River - West Fork Coal Creek | 166 | 1993 | 61.482, -151.633 | 95 | 0.737 | 0.263 | 0 | 0 | 0 | 0 | 0 |
| 25 | | Little Jack Creek | 170 | 9/06/2006 | 60.58, -152.362 | 95 | 0.821 | 0.147 | 0 | 0 | 0 | 0 | 0.032 |

| | | | | | | | | | | | | | |
|---|---|---|---|---|---|---|---|---|---|---|---|---|---|
| 26 | Yentna Drainage | Puntilla Lake | 177 | 9/06/2006 | 62.087, -152.733 | 95 | 0.66 | 0.34 | 0 | 0 | 0 | 0 | 0 |
| 27 | Susitna Drainage | Larson Lake | 185 | 09/1993 | 62.355, -149.866 | 95 | 0.581 | 0.384 | 0 | 0 | 0.035 | 0 | 0 |
| 28 | Knik Arm | Six Mile Creek | 198 | 1997 | 61.294, -149.835 | 95 | 0.659 | 0.341 | 0 | 0 | 0 | 0 | 0 |
| 29 | Cook Inlet - Northeast | Bishop Creek | 199 | 1993 | 60.784, -151.079 | 95 | 0.736 | 0.264 | 0 | 0 | 0 | 0 | 0 |
| 30 | Kenai Drainage | Hidden Creek | 202 | 7/29/1993 | 60.466, -150.209 | 95 | 0.074 | 0.663 | 0 | 0 | 0 | 0 | 0.263 |
| 31 | | Upper Russian River weir - late | 208 | 8/02/1993 | 60.486, -150.003 | 95 | 0.105 | 0.011 | 0 | 0 | 0 | 0 | 0.884 |
| 32 | Kasilof Drainage | Bear Creek | 219 | 8/10/1993 | 60.213, -150.806 | 95 | 0.137 | 0.842 | 0 | 0 | 0 | 0 | 0.021 |
| 33 | Outer Gulf Coast | Delight River | 225 | 1993 | 59.538, -150.35 | 71 | 0.408 | 0.592 | 0 | 0 | 0 | 0 | 0 |
| 34 | Prince William Sound | Coghill Lake | 227 | 09/1991 | 61.08, -147.849 | 96 | 0.2 | 0.779 | 0 | 0 | 0 | 0 | 0.021 |
| 35 | Copper River Drainage | Bering Lake | 228 | 7/12/1991 | 60.28, -144.304 | 95 | 0.222 | 0.578 | 0 | 0 | 0 | 0 | 0.2 |
| 36 | | Eyak Lake - South Beaches | 230 | 06/1991 | 60.544, -145.679 | 39 | 0.051 | 0.872 | 0 | 0 | 0 | 0 | 0.077 |
| 37 | | St. Anne Creek | 232 | 7/15/2005 | 61.826, -146.052 | 95 | 0.065 | 0.935 | 0 | 0 | 0 | 0 | 0 |
| 38 | | Tanada Creek weir | 233 | 8/21/2005 | 62.596, -143.718 | 95 | 0 | 0.83 | 0 | 0 | 0 | 0 | 0.17 |
| 39 | Yakutat | East Alsek | 234 | 10/15/2000 | 59.153, -138.584 | 96 | 0.149 | 0.298 | 0 | 0 | 0 | 0 | 0.553 |
| 40 | Trans-Boundary | Alsek River - Klukshu River Weir - late | 235 | 8/23/2006 | 60.163, -137.225 | 95 | 0.158 | 0.674 | 0 | 0 | 0 | 0 | 0.168 |
| 41 | | Taku River - Little Tatsamenie | 238 | 9/11/1991 | 58.456, -132.396 | 81 | 0 | 0.485 | 0 | 0 | 0 | 0 | 0.515 |
| 42 | | Stikine River - Iskut River | 242 | 1986 | 56.75, -131.783 | 84 | 0.051 | 0.859 | 0 | 0 | 0 | 0 | 0.09 |
| 43 | Northern Southeast | Sitkoh Lake | 252 | 9/26/2003 | 57.514, -135.045 | 95 | 0 | 0.606 | 0 | 0 | 0 | 0 | 0.394 |
| 44 | | Falls Lake | 253 | 9/02/2003 | 56.825, -134.692 | 95 | 0.263 | 0.021 | 0 | 0 | 0 | 0 | 0.716 |
| 45 | Southern Southeast | Luck Lake - P.O.W. Island | 263 | 9/10/2004 | 55.953, -132.759 | 95 | 0.18 | 0.101 | 0 | 0 | 0 | 0 | 0.719 |
| 46 | | Hetta Lake | 272 | 10/01/2003 | 55.171, -132.568 | 93 | 0.012 | 0.554 | 0 | 0 | 0.024 | 0 | 0.41 |
| 47 | Nass River | Hanna Creek | 279 | 9/03/2006 | 56.083, -129.3 | 95 | 0.333 | 0.129 | 0 | 0 | 0 | 0 | 0.538 |
| 48 | Skeena River | Alastair Lake | 283 | 9/14/2006 | 54.1, -129.183 | 86 | 0.153 | 0.259 | 0 | 0 | 0 | 0 | 0.588 |
| 49 | | Johanson Lake - Sustut | 294 | 2006 | 56.583, -126.183 | 95 | 0.484 | 0.105 | 0 | 0 | 0 | 0 | 0.411 |
| 50 | Central Coast | Kitlope Lake | 297 | 8/03/2006 | 53.117, -127.783 | 95 | 0.457 | 0.468 | 0 | 0 | 0.011 | 0 | 0.064 |
| 51 | Queen Charlotte Is. | Naden River | 298 | 1995 | 53.947, -132.667 | 95 | 0 | 0.149 | 0 | 0 | 0 | 0 | 0.851 |
| 52 | Fraser River | Lower Horsefly River1 | 300 | 9/12/2001 | 52.467, -121.383 | 190 | 0 | 0.878 | 0 | 0 | 0 | 0 | 0.122 |
| 53 | | Weaver Creek | 302 | 2001 | 49.317, -121.883 | 95 | 0 | 0.372 | 0 | 0 | 0.106 | 0 | 0.521 |
| 54 | Washington/Idaho | Baker Lake | 303 | 5/16/1996 | 48.548, -121.741 | 97 | 0 | 0.853 | 0 | 0 | 0.032 | 0 | 0.116 |
| 55 | | Cedar River | 304 | 10/26/1994 | 47.5, -122.216 | 96 | 0.094 | 0.875 | 0 | 0 | 0 | 0 | 0.031 |
